# α-Helical peptidic scaffolds to target α-synuclein pathogenic species with high affinity and selectivity

**DOI:** 10.1101/2021.03.12.435074

**Authors:** Jaime Santos, Pablo Gracia, Susanna Navarro, Samuel Peña-Diaz, Jordi Pujols, Nunilo Cremades, Irantzu Pallarès, Salvador Ventura

## Abstract

α-Synuclein aggregation is a key driver of neurodegeneration in Parkinson’s disease and related syndromes. Accordingly, obtaining a molecule that targets α-synuclein pathogenic assemblies with high affinity and selectivity is a long-pursued objective. Here, we have exploited the biophysical properties of toxic oligomers and amyloid fibrils to identify a family of α-helical peptides that bind selectively to these α-synuclein species with low nanomolar affinity, without interfering with the monomeric functional protein. This activity is translated into an unprecedented anti-aggregation potency and the ability to abrogate the oligomers toxicity. With a structure-function relationship in hand, we identified a human peptide expressed in the brain and in the gastrointestinal tract with exceptional binding, antiaggregation, and detoxifying properties, which suggests it might play a protective role against synucleinopathies. The chemical entities we describe here represent a new therapeutic paradigm and are promising tools to assist diagnosis by selectively detecting α-synuclein pathogenic species in biofluids.

## Introduction

α-Synuclein (αS) is a 140 amino acid protein whose aggregation into amyloid fibrils in a subset of neuronal and glial cells lies behind the onset of a group of progressive and, ultimately, fatal neurodegenerative disorders, including Parkinson’s disease (PD)^1–4^, that are collectively referred to as synucleinopathies. A causative link between αS and disease is supported by the discoveries that multiplications and missense mutations in SNCA, the αS gene, cause dominantly-inherited familial forms of PD^5^.

Interfering with αS amyloid formation and abrogating the associated toxicity is considered a promising therapeutic strategy for synucleinopathies^6–8^. However, the design of molecular entities that target specific αS pathogenic assemblies is challenging because of the heterogeneous, dynamic, and transient nature of these species. High-throughput screening initiatives have rendered promising αS aggregation inhibitors^9, 11^. However, these selection procedures are blind to the ligand mechanism of action. In the absence of a structure-activity relationship (SAR), it is difficult to evolve the affinity and specificity of the identified hits to generate drugs that can reach the clinics. The lack of specific and sensitive molecules to detect the pathogenic forms of αS also constrains the early diagnosis of these diseases.

The *in vitro* aggregation of αS displays a sigmoidal growth profile, suggesting that it follows a nucleation-polymerization mechanism^12^, where soluble αS undergoes a nucleation process that produces oligomers able to grow through further monomer addition to form insoluble amyloid fibrils. Oligomeric forms of αS have been detected in the brains and other tissues of patients suffering from PD, and growing evidence suggests that they constitute the primary pathogenic agents accounting for the gain-of-toxicity associated with αS aggregation, whereas both oligomers and fibrils would be responsible for pathology dissemination in the brain^2,13–15^. We have recently identified the sequential population of two conformationally distinct types of oligomers during αS *in vitro* fibrillation. The initial non-toxic disordered oligomers, named as type A oligomers, undergo a structural reorganization to form more stable and compact β-sheet enriched, and proteinase-K resistant species that exhibit intrinsic cytotoxicity, named as type B oligomers^16^. Stable, trapped analogous of these two well-defined types of transient oligomers (referred to as type A* and type B* oligomers, where the star refers to the kinetically trapped nature of these isolated oligomeric forms) have been isolated and characterized in detail^13,16^ and, therefore, constitute important tools for the development of specific therapeutic and diagnostic strategies.

In this work, we have exploited our recent advances in the understanding of the structural determinants of toxicity of αS oligomers to rationally identify novel molecules able to target the pathogenic species of aS. By using a time-resolved single-particle fluorescence approach we demonstrate that short, amphipathic and cationic α-helical peptides bind toxic oligomers and fibrils with unprecedented specificity and affinity, resulting in the substoichiometric inhibition of αS aggregation and abrogation of oligomer toxicity in neuronal cell models. We then used a protein engineering approach to dissect the molecular determinants accounting for this interaction, which has allowed us to identify a human peptide, constitutively expressed in the brain and gastrointestinal tract, that binds with low nanomolar affinity and high specificity to αS pathogenic species, thus suppressing the aggregation cascade and its associated neurotoxicity, which suggests it might play a natural protective role against synucleinopathies.

In summary, here we describe the rational identification and characterization of a family of highly potent and αS species-specific peptidic ligands able to bind selectively to αS toxic species and abrogate their toxic effects in neuronal cells. This discovery opens previously unexplored avenues for the diagnosis and/or therapeutics of PD and related disorders.

## Results

### Identification of an αS species-specific peptide ligand

We rationalized that the particular properties of the four main αS conformers identified during αS amyloid aggregation, namely monomers, non-toxic (type A/A*) oligomers, toxic (type B/B*) oligomers and fibrils, could be exploited to identify a selective ligand for the main species responsible for induction and propagation of toxicity, which are currently believed to be type B-like oligomers and amyloid fibrils, respectively^17^. In **Figure 1a**, we illustrate the dissection of the differential traits of αS forms. Type B-like oligomers and amyloid fibrils share two features: (i) they expose relatively large lipophilic clusters to the solvent. These hydrophobic surfaces induce cellular toxicity and drive subsequent fibrillation^16,18, 20^. (ii) They possess a high anionic character, at neutral pH as a result of the stacking of αS monomers (net charge −9) with the highly negatively charged C-terminal regions exposed to the solvent. While the solvent exposure of hydrophobic surfaces seems to be a general feature of toxic pre-fibrillar oligomers^13,21^, the combination of high exposed hydrophobicity and negative charge is likely unique to these two pathogenic αS assemblies. Thus, we hypothesized that hydrophobic patches embedded in an anionic environment might delineate a diffuse, but physicochemically-defined, binding surface in these two types of αS aggregates for a complementary molecule; ideally an amphipathic and cationic entity. A short α-helical peptide might provide a structurally stable scaffold to merge both features.

**Figure 1.**
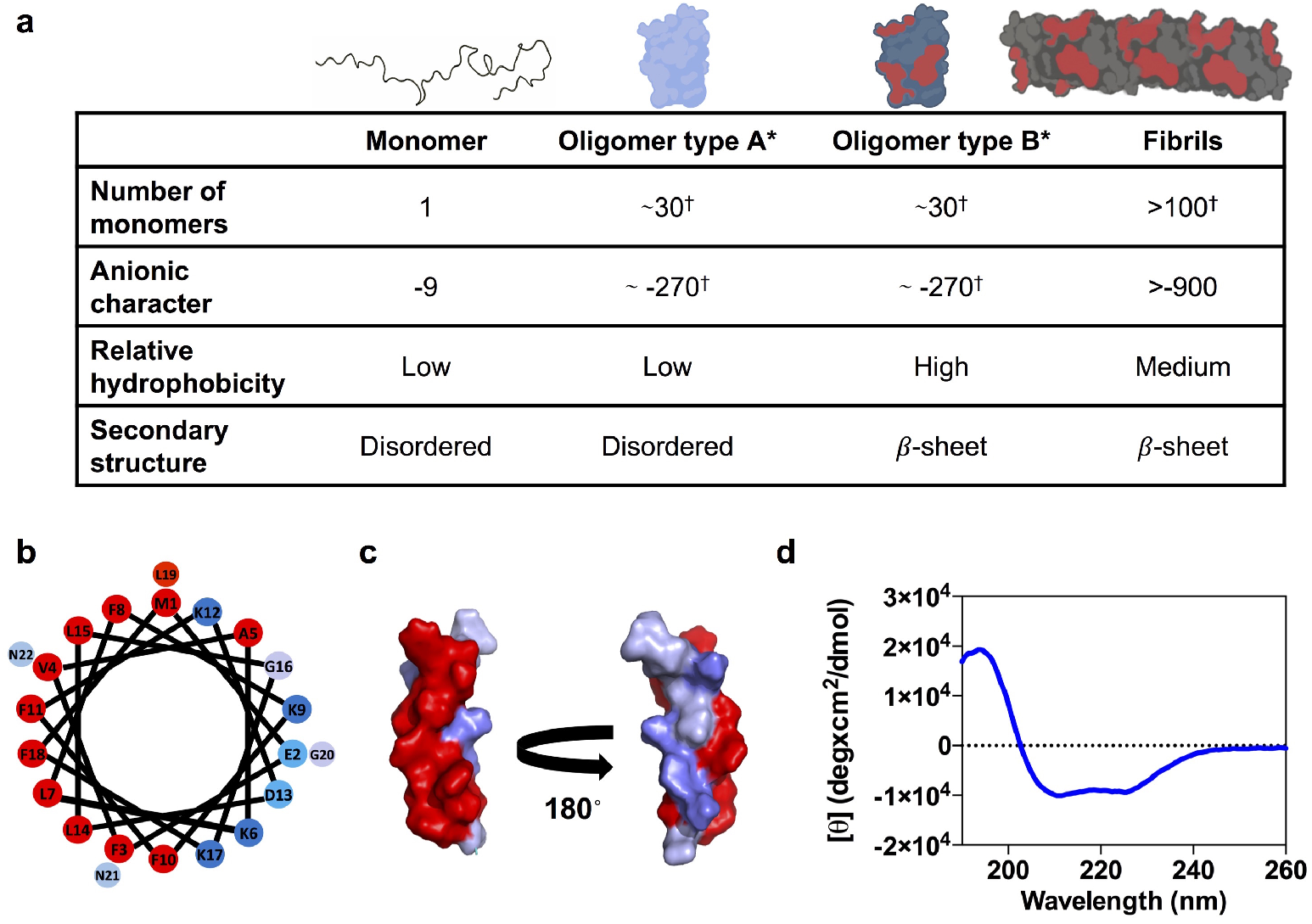
Rational identification of a peptide ligand for αS pathogenic species. (**a**) Main molecular features of the four isolated αS species. Values with a dagger (†) represent extrapolations based on the average number of monomers in each specie. (**b**) Helical wheel projection of PSMα3 sequence (red, hydrophobic residues; blue pallet, hydrophilic residues depending on their character). (**c**) Surface representation of the three-dimensional structure of PSMα3 with hydrophobic residues in red and hydrophilic residues in blue. (**d**) Far-UV circular dichroism spectra of PSMα3.

Short α-helical peptides tend to be only marginally stable in solution. Therefore, the *de novo* design of an amphipathic and cationic helical chassis that retains a stable secondary structure under physiological conditions is not trivial^22,23^. Instead, we decided to identify a naturally occurring peptide bearing our desired properties, assuming that evolution would have selected for a stable α-helix fold. This is especially true for secreted peptides, which should remain folded in a harsh extracellular environment. Following this rationale, we selected PSMα3 as a candidate peptide, since, theoretically, it fulfilled all our requirements. PSMα3 is a 22 residues bacterial extracellular peptide that has been shown to remain in an α-helical conformation for weeks^24^. It has a net charge of +2, a mean hydrophobicity *(H)* of 0.54, an α-helical hydrophobic moment (μ_*H*_) of 0.56, and the helical wheel plot evidences its amphipathic character (**Figure 1b-c**). The far-UV circular dichroism (CD) spectrum of PSMα3 confirms that it folds into an α-helix under our assay conditions (**Figure 1d**).

We engineered PSMα3 to obtain a negative control peptide in which the formation of an α-helix is strongly disfavored. This will disrupt the peptide amphipathic character and, theoretically, abolish binding to αS type B* oligomers and amyloid fibrils. After a computational proline scanning of PSMα3 using the AGADIR algorithm^25^ (**Supplementary Figure 1a**), we selected the K9P and F11P mutations, as they have a significant impact in helical propensity and map to opposite faces of the α-helix (**Supplementary Figure 1b-c)**. The characterization of the secondary structure of the K9P-F11P PSMα3 peptide (further referred to as disrupted PSMα3 or dPSMα3) in solution by CD confirmed the disruption of the α-helix fold (**Supplementary Figure 1d**). Thus, dPSMα3 constitutes a suitable negative control for further studies, as it keeps a sequence identity of 91 % with PSMα3, but lacks its amphipathic character, a feature that we propose is key for the specific binding to the αS toxic species.

### Selective interaction of PSMα3 with αS pathogenic species

We then addressed the interaction of PSMα3 and dPSMα3 with the four major αS species generated during the process of amyloid aggregation. As multiple peptides are expected to bind multiple αS molecules in the aggregated states, the binding process can only be well described if both the stoichiometry of the complex and the affinity of the peptide for the αS molecules in a particular conformer is known. In order to obtain good estimates of both parameters, we exploited the power of dual-color fluorescence cross-correlation spectroscopy (dcFCCS), a time-resolved fluorescence fluctuation technique that allows the direct observation of co-diffusing fluorescent species arising from interactions between differently labeled molecules or assemblies in solution^26,27^. To this end, αS species were cys-labeled with maleimide-AlexaFluor488 (AF488), with each αS molecule of the different species containing one fluorophore at position 122, and the peptides were cys-labelled with maleimide-Atto647N at the N-terminus (see Methods section for details). Simultaneously, we assessed complex formation by single-particle fluorescence spectroscopy (SPF) analysis, including Förster Resonance Energy Transfer (FRET) and donor/acceptor stoichiometry (*S*), in order to validate and complement the dcFCCS approach. These approaches allow us to monitor distinct individual species simultaneously by avoiding measurements of ensemble averages (**Supplementary Figure 2**) and have been previously used for the characterization of αS aggregation pathways and the study of αS interactors^28,30^.

We first assessed the binding of PSMα3 to monomeric αS and found by dcFCCS analysis that these molecules were unable to interact when mixed in an equimolar ratio (15 nM of each molecule) (**Figure 2a**), as reflected by a flat cross-correlation curve comparable to that of the negative control of cross-correlation (**Supplementary Figure 3**). We then analyzed the interaction of PSMα3 with the different αS aggregated species by means of dcFCCS at approximately equimolar ratios of peptide and αS molecules (1-5 nM). Due to the different stability of the various aggregated species upon single-molecule dilution and their differential adsorption to the surface of the coverslips, the total αS concentration of each aggregated sample was adjusted between 1 and 5 nM (mass concentration) so that the frequency of events in the diffusion single-particle measurements was very similar between the various aggregated samples. Under these conditions, a marginal cross-correlation amplitude was observed for the type A* oligomers (**Figure 2b**), whereas a clear cross-correlation was obtained for the interaction of the peptide with both type B* oligomers and fibrils (**Figure 2c and d**, respectively), already indicating a stark difference in the binding ability of the peptide to the different aggregates. Consistently, a high number of FRET events were detected by singleparticle burst-wise analysis, thus validating the direct interaction of the peptide with these two αS species. In contrast, the same analysis yielded either a statistically insignificant number of FRET events or none at all for the interaction between PSMα3 and αS type A* oligomers and monomers, respectively (**Supplementary Figure 4a-c**). These results offer a single-particle understanding of a complex binding scenario and further reinforce the observations derived from dcFCCS experiments.

**Figure 2:**
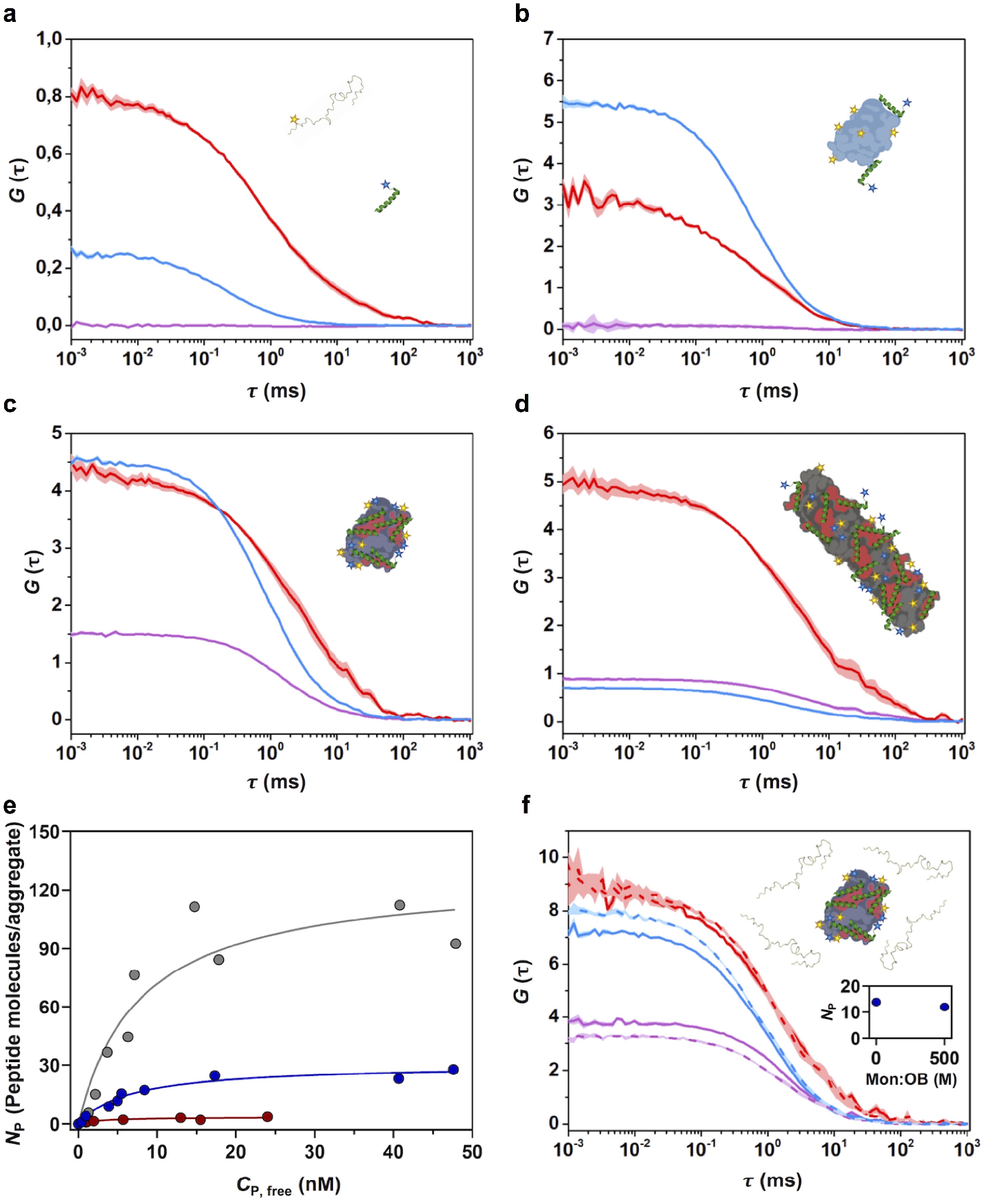
Interaction of PSMα3 with different αS species by FCCS. (**a-d**) Representative auto-correlation curves for αS (blue line) and PSMα3 (red line) and cross-correlation curves for interacting molecules (purple line). The amplitude (*G*) error is shown as faint colored area for the corresponding correlation curves. Samples contained (**a**) ~15 nM αS monomer and ~15 nM PSMα3, (**b**) 1 nM type A* and ~5 nM PSMα3, (**c**) 1nM type B* oligomers and ~5 nM PSMα3 or (**d**) ~5 nM sonicated fibrils and ~5 nM PSMα3. (**e**) Titration binding curves for the interaction of PSMα3 with type A* oligomers (red circles), type B* oligomers (blue circles) or sonicated fibrils (grey circles) obtained by FCCS experiments, showing their corresponding analysis assuming a model of *n* identical and independent binding sites (referred in equation 7 as N_max_) per αS aggregated species (solid lines). (**f**) Auto-correlation curves (αS in blue, PSMα3 peptide in red) and cross-correlation curve for the interacting molecules (in purple) obtained in samples containing ~1 nM αS type B* oligomers and ~2 nM PSMα3 in the absence (solid lines) or presence (dashed lines) of a 500-molar excess of unlabeled monomer with respect to the particle concentration of oligomers. The inset shows the number of bound peptides (*N*_P_) per aggregate in both conditions.

We next performed a titration experiment for analyzing the binding of the peptide to the three αS aggregated species. For this, we developed a model-independent saturation binding curve, based on the theoretical framework previously developed by Kruger and coworkers^31^. This analysis allowed us to quantify the number of peptide molecules bound to each αS species (*N*_p_) as a function of the concentration of unbound peptide (*C*_p,Free_) (**Figure 2e**). Using a simplistic Langmuir isotherm model, we estimated the single-state dissociation constant (*K_D_*) of the interactions and the average maximum number of peptide binding sites (*N*_max_) in each type of αS species. Interestingly, while the *K_D_* values for the peptide-aS interaction obtained for type A* oligomers, type B* oligomers and fibrils are very similar, in all cases in the very low nM range (3.07 nM, 6.67 nM and 7.8 nM, respectively), the average maximum number of peptides per aggregate varies remarkably, being 3, 30 and 120, respectively. This indicates that the main difference between the three aggregated species in terms of PSMα3 interaction is the number of binding sites per aggregate rather than the affinity of the peptide for them. It is remarkable that the average maximum number of binding sites obtained for the type B* oligomers and fibrils nearly matches the average number of αS molecules per aggregate species (19, 33 and 107 for type A*, type B* oligomers and fibrils, respectively, estimated by comparing the molecular brightness of the aggregated species to that of the αS monomer, not shown), while for the type A* oligomers represents only one sixth of the average αS molecules in this type of aggregate. This data is in agreement with the single-particle fluorescence analysis obtained for the different complexes, where only very few FRET events were observed for the binding of PSMα3 to type A* oligomers, in contrast to the numerous FRET events with a defined FRET efficiency (*E*) distribution observed for the binding to type B* oligomers and fibrils (**Supplementary Figure 4**). Note that a 500-fold excess of unlabeled monomeric αS does not interfere with the binding of PSMα3 to type B* oligomers (**Figure 2f**) which indicates a high specificity towards aggregated species and negligible monomer binding. Together, the dcFCCS and single-particle fluorescence data demonstrate that PSMα3 is a high affinity αS species-specific ligand, with affinities in the low nanomolar range, and with a remarkable avidity and selectivity for the type B* oligomers and fibrils. Interestingly, when we analyzed the binding of the PSMα3 analogous, but disordered peptide, dPSMα3, we could not detect any interaction with any of the four αS species (**Supplementary Figure 5**), indicating that an amphipathic distribution of the peptide residues, achieved through an α-helical conformation, is a requirement for the interaction.

Overall, our dcFCCS-derived binding analysis indicates that PSMα3 binds with low nanomolar affinity to all αS aggregated species. The degree of binding is limited by the number of available interaction sites, which is likely associated with the extent of solvent-exposed hydrophobic surface per aggregate, which depends on both the size and the lipophilicity of the aggregate, in agreement with our initial reasoning.

### PSMα3 inhibits αS amyloid aggregation

We hypothesized that the high affinity and number of binding sites of type B* oligomers for PSMα3 might result in the partial or full coverage of the surface of these assemblies and likely their structurally homologous type B oligomers, thus preventing their progression to fibrils during the αS amyloid aggregation process. To assess if this was the case, we set up *in vitro* αS aggregation reactions in the absence and presence of an equimolar concentration of PSMα3 (70 μM) and followed its progression by monitoring the increase in thioflavin-T (Th-T) fluorescence. After 32 hours of incubation, a ~90 % decrease in Th-T fluorescence emission, relative to the untreated sample, was observed in the presence of PSMα3, suggesting that the peptide acts as a potent inhibitor of αS amyloid aggregation (**Figure 3a**). Transmission electron microscopy (TEM) images confirmed that samples incubated with PSMα3 contained very few fibrils per field, in comparison to untreated samples (**Figure 3b).** The inhibitory activity of PSMα3 was concentration-dependent and significant even at a substoichiometric 20:1 ratio (αS:PSMα3) (**Figure 3c**), in agreement with its selective binding to low populated oligomeric species formed during the early stages of αS amyloid aggregation, in line with the time-resolved fluorescence spectroscopy data. The observation that dPSMα3 exhibited a negligible anti-aggregative activity (**Supplementary Figure 6**) reinforces the connection between the binding of the amphipathic (hydrophobic/cationic) helical peptide to αS oligomers and its potent amyloid inhibition activity.

**Figure 3.**
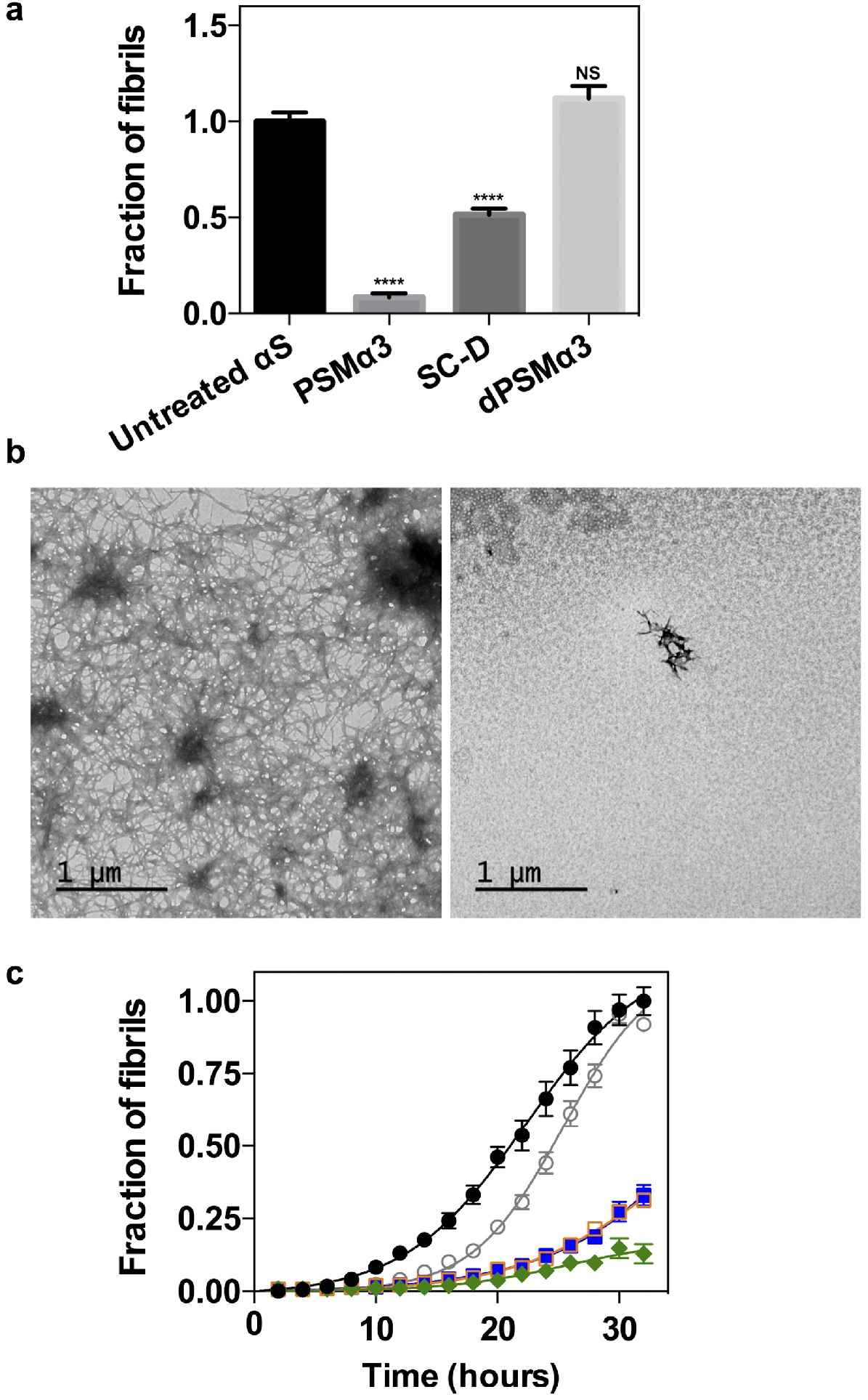
Effect of PSMα3 on *in vitro* αS amyloid fibrillation. (**a**) Inhibition of αS amyloid aggregation as measured by Th-T fluorescence after 32 h incubation in the presence of equimolar concentrations of PSMα3, SynuClean-D (SC-D), and dPSMα3. **** p-value < 0.0001 relative to untreated αS (**b**) TEM micrographs of αS aggregated for 32 h in the absence (left) and presence of an equimolar concentration of PSMα3 (right). (**c**) Aggregation kinetics of 70 μM αS and titration of the inhibitory activity of PSMα3 at different concentrations: 35 μM (green), 14 μM (orange), 7 μM (blue), 3.5 μM (gray) and in the absence of PSMα3 (black). Error bars represent the standard error of the mean (SEM).

Remarkably, PSMα3 is a better inhibitor of αS amyloid formation than SynuClean-D (**Figure 3a**), a small molecule with high neuroprotective activity in *Caenorhabditis elegans* models of PD that we have discovered recently^9^. Indeed, a survey of the literature indicates that the inhibition potency of PSMα3 exceeds that reported for most non-covalent small molecules, including the diphenylpyrazole Anle138b, already in clinical trials^10^.

### PSMα3 protects cells from αS oligomer-induced toxicity

As it occurs with toxic oligomers from other amyloidogenic systems, the toxicity of type B* oligomers relies on their ability to interact and disrupt cellular membranes^13^. In αS this activity is encoded in two of their characteristic structural elements^13^: (i) an exposed N-terminal region that acts as the initial anchor to the membrane surface, similarly as with the monomeric functional form of the protein, and (ii) a β-sheet core, composed primarily by the central region of the protein, with significant hydrophobic surface exposed to the solvent that then inserts into the lipid bilayer causing major perturbations. The highly negatively charged C-terminal region of the protein remains disordered without significant interactions with the membrane.

We hypothesized that the binding of PSMα3 to type B* oligomers, likely mediated by the solvent exposed hydrophobic regions of the β-sheet core together with the proximal negatively charged C-terminal regions, would block their exposed lipophilic elements, thus decreasing its ability to insert to and perturb the membrane bilayer and, therefore, to induce cellular toxicity. We treated human SH-SY5Y neuroblastoma cells with 10 μM of oligomers and observed that, as previously reported^32^, they possess a high affinity for cellular membranes (**Figure 4a-b**). When the oligomers were preincubated with an equimolar concentration of PSMα3, we observed a ~60 % reduction in the amount of αS bound to cells relative to untreated oligomers, indicating that PSMα3 binding to type B* oligomers directly affects the binding of the oligomers to cellular membranes.

**Figure 4.**
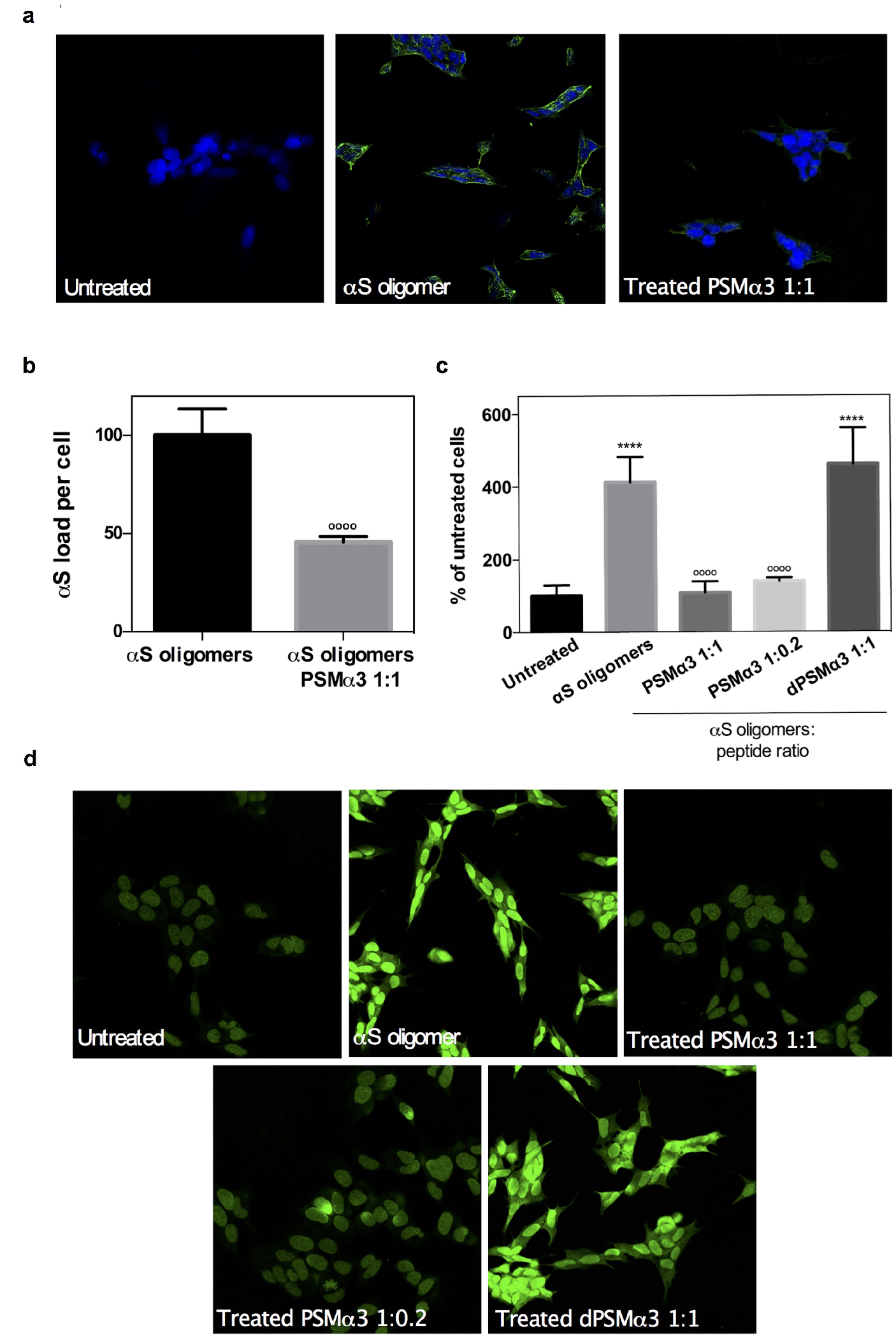
Suppression of the toxicity of αS oligomers in neuroblastoma cells. (**a**) Representative confocal images showing the αS load per cell after the treatment with 10 μM of type B* oligomers pretreated with an equimolar concentration of PSMα3. (**b**) Quantification of the αS load per cell. Error bars represent the standard deviation (SD). **** p-value < 0.0001 relative to untreated cells. ^oooo^ p-value < 0.0001 relative to cells treated with αS type-B* oligomers. (**c**) Quantification of the levels of intracellular ROS of SH-SY5Y cells incubated with 10 μM of type B* oligomers preincubated with different concentrations of PSMα3 and dPSMα3. Error bars represent the standard error of the mean (SEM). ^oooo^ p-value < 0.0001 relative to cells treated with αS type B* oligomers. (**d**) Representative confocal images of the analysis of panel c.

One of the earliest effects of type B oligomer-mediated membrane perturbation is the substantial increase in the levels of intracellular reactive oxygen species (ROS)^21^, which in turn elicits mitochondrial dysfunction^33^. We assessed if the blockage of the oligomer regions involved in membrane perturbation by PSMα3 binding could protect membrane integrity and therefore prevent its associated increase in intracellular ROS levels. Treatment of neuroblastoma cells with 10 μM of oligomers induced a drastic increase in ROS levels (**Figure 4c-d**). However, when these oligomers were preincubated with equimolar (1:1) and substoichiometric (1:0.2) concentrations of PSMα3, the ROS levels of treated cells approached those of healthy, untreated, cells, indicating that PSMα3 protects against oligomers induced damage. This detoxifying activity seems to be associated with the particular structural and physicochemical properties of this peptide since treatment with equimolar concentrations of dPSMα3 failed to exert any protective effect.

### Dissection of PSMα3 aggregation-inhibitory determinants

To this point, we have assigned the αS binding, anti-aggregation and cytoprotective properties of PSMα3 to its helical, amphipathic and cationic character. To confirm that this is the case, we reverse-engineered PSMα3 into a non-natural peptide scaffold with low sequence complexity that keeps its critical properties. We employed a set of bioinformatics tools to predict the helical propensity, helical hydrophobic moment, and thermodynamic stability of our successive designs using AGADIR^25^, HELIQUEST^34^, and FOLDX^35^, respectively. Data regarding those predictions are displayed in **Supplementary Table 1**. Then, we evaluated the anti-aggregative properties of these new molecules, under the assumption that the inhibitory capacity is connected with the oligomer-peptide interaction affinity.

A first requirement for binding is a continuous hydrophobic face to interact with the surface of oligomers or fibrils. In our view, the specific sequence of this helical side would be irrelevant, as long as it keeps its lipophilic character. To demonstrate that this assumption is correct, we mutated all the residues in the hydrophobic face of the PSMα3 α-helix to leucine (All_Leu), generating an amphipathic peptide devoid of any sequence diversity in this side. Simultaneously, we designed a variant of All-Leu devoid of the last three C-terminal residues (All_Leu19) since they are not part of the α-helix, and thus they are not expected to contribute significantly to the binding. Both redesigned peptides folded into α-helices and retained the inhibitory activity, with a potency that approaches that of PSMα3 (**Figure 5a-b**). Thus, we concluded that it is the generic hydrophobic character of the helical face and not its sequence or composition that is relevant for the binding.

**Figure 5.**
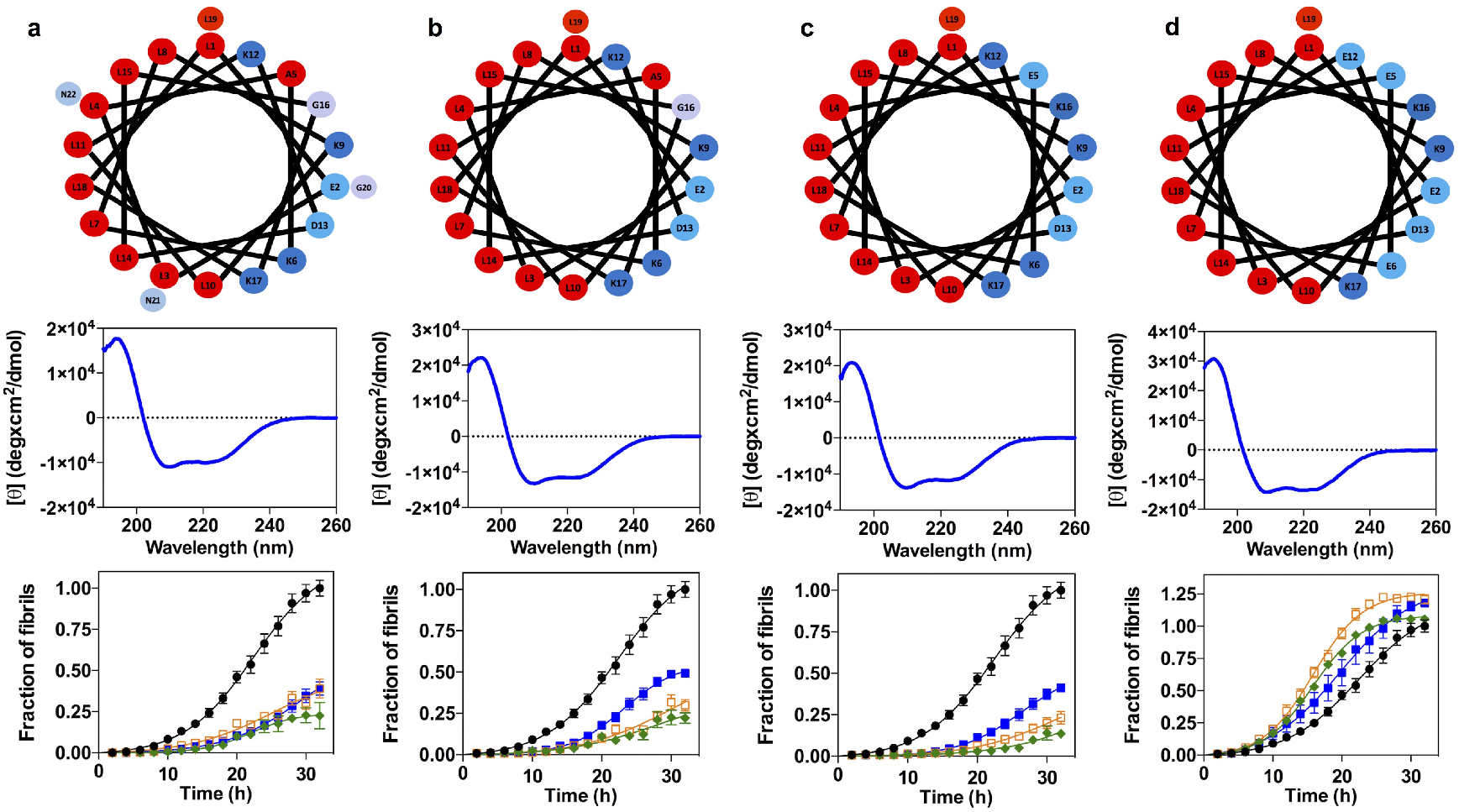
Redesign of PSMα3 variants to dissect the molecular determinants of the anti-aggregative activity. (**a,b,c,d**) Helical wheel (up), circular dichroism spectra (mid) and titration of the inhibitory activity of PSMα3 at different concentrations (down): 35 μM (green), 14 μM (orange), 7 μM (blue) and in absence of PSMα3 variants (black). Variants: All_Leu (**a**), All_Leu19 (**b**), Scaffold_19 (**c**) and Anionic_scaffold (**d**). Error bars represent the standard error of the mean (SEM).

Next, to further reduce the peptide sequence complexity, we redesigned the hydrophilic face in such a way that it only contained ionizable residues. We designed a new variant (Scaffold_19) with only four different amino acids (Leu, Asp, Glu, and Lys) by introducing two point-mutations (A5E_G16K) in the All_Leu19 peptide. We decided to maintain the peptide net charge by introducing residues with opposed charges. Thus, Scaffold_19 only has Leu in the hydrophobic face and charged residues in the hydrophilic one. This variant folded into an α-helix and showed the same anti-aggregation activity than the parental variant (**Figure 5c**).

The simplicity of Scaffold_19 allowed us to redesign the net charge of the peptide, to validate the other physicochemical property theoretically contributing to binding: a net positive charge. We generated a peptide with a net charge of −2 by introducing two charge-reversing mutations (K6E_K12E). This anionic peptide (Anionic_scaffold) folds into an α-helix and has a helical hydrophobic moment (μ_*H*_) of 0.65, indicative of an amphipathic nature, but does not inhibit αS amyloid aggregation, confirming that a cationic character in the hydrophilic face is a requirement for binding (**Figure 5d**).

Overall, we succeeded in dissecting the peptide features responsible for aggregation inhibition. In the process, we generated a short peptide with low sequence complexity whose inhibitory activity does not stem from the primary sequence, but instead from a defined spatial distribution of two physicochemical traits. Scaffold_19 constitutes a representative of a generic family of peptides with potential therapeutic applications.

### LL-37 inhibits αS aggregation and oligomer toxicity

Once we elucidated the determinants of this novel mechanism of αS amyloid inhibition, we wondered if this activity could also be encoded in natural human peptides. First, we screened the EROP-Moscow oligopeptide database^36^ for human cationic peptides longer than 10 residues (? 3 helical turns), obtaining 287 hits. Next, we run AGADIR on them to exclude peptides with a low helical propensity, which resulted in 25 peptides, from which only 9 peptides were predicted to have at least a partial amphipathic character, according to their helical hydrophobic moment (μ_*H*_) (**Supplementary Table 2**). Then we screened the literature for candidates whose tissue distribution overlapped with that of αS and selected LL-37, the only human member of the cathelicidin family of antimicrobial peptides, for its further characterization. LL-37 is a 37-residue peptide resulting from a post-translational cleavage at the C-terminus of cathelicidin hCAP18^37^. This peptide is constitutively expressed in the brain and the gastrointestinal tract; its presence in both tissues is engaging, as the brain-gut axis connection is gaining momentum in PD^38–40^.

First, we confirmed that LL-37 adopts an α-helical conformation under our assay conditions. With α-helical hydrophobic moment (μ_*H*_) of 0.52, the helical-wheel projection and the available 3D-structures^34^ indicate that this α-helix would be both cationic and amphipathic (**Figure 6a-b**). Then, we titrated the anti-aggregative activity of LL-37, confirming that it suppresses αS amyloid formation at substoichiometric concentrations (**Figure 6c**). Next, we labeled LL-37 with maleimide-Atto647N at a single engineered cysteine at the N-terminus, and we performed time-resolved dual-color fluorescence spectroscopy experiments as described previously for PSMα3. Both dcFCCS and spFRET (**Figure 6d-f and Supplementary Figure 7**) reported a strong binding to type B* oligomers and fibrils, a weak interaction with type-A* oligomers and the absence of any interaction with the αS monomer, indicating that LL-37 and PSMα3 share a very similar binding mechanism (**Figure 6g**), with LL-37 displaying slightly higher affinities than PSMα3 (*K_D_* = 3.62 nM for type B* oligomers, *K_D_* = 5.14 nM for sonicated fibrils and *K_D_* = 1.92 nM for type A* oligomers), and significant higher number of binding sites in type B* oligomers (*N*_max_ = 64) and sonicated fibrils (*N*_max_ = 181), while remain the same as PSMα3 for the number of binding sites in the type A* oligomers (*N*_max_ = 3). Consistent with the LL-37 ability to bind type B* oligomers with high affinity, the preincubation of these toxic species with the human peptide at an equimolar concentration completely abolished the production of ROS in neuroblastoma cells (**Figure 6h-i**). LL-37 is not related in sequence to PSMα3 or Scaffold_19, but the three peptides share the same structural and physicochemical traits. This confirms that a linear combination of these properties suffices to identify, and potentially design, potent inhibitors of αS aggregation.

**Figure 6.**
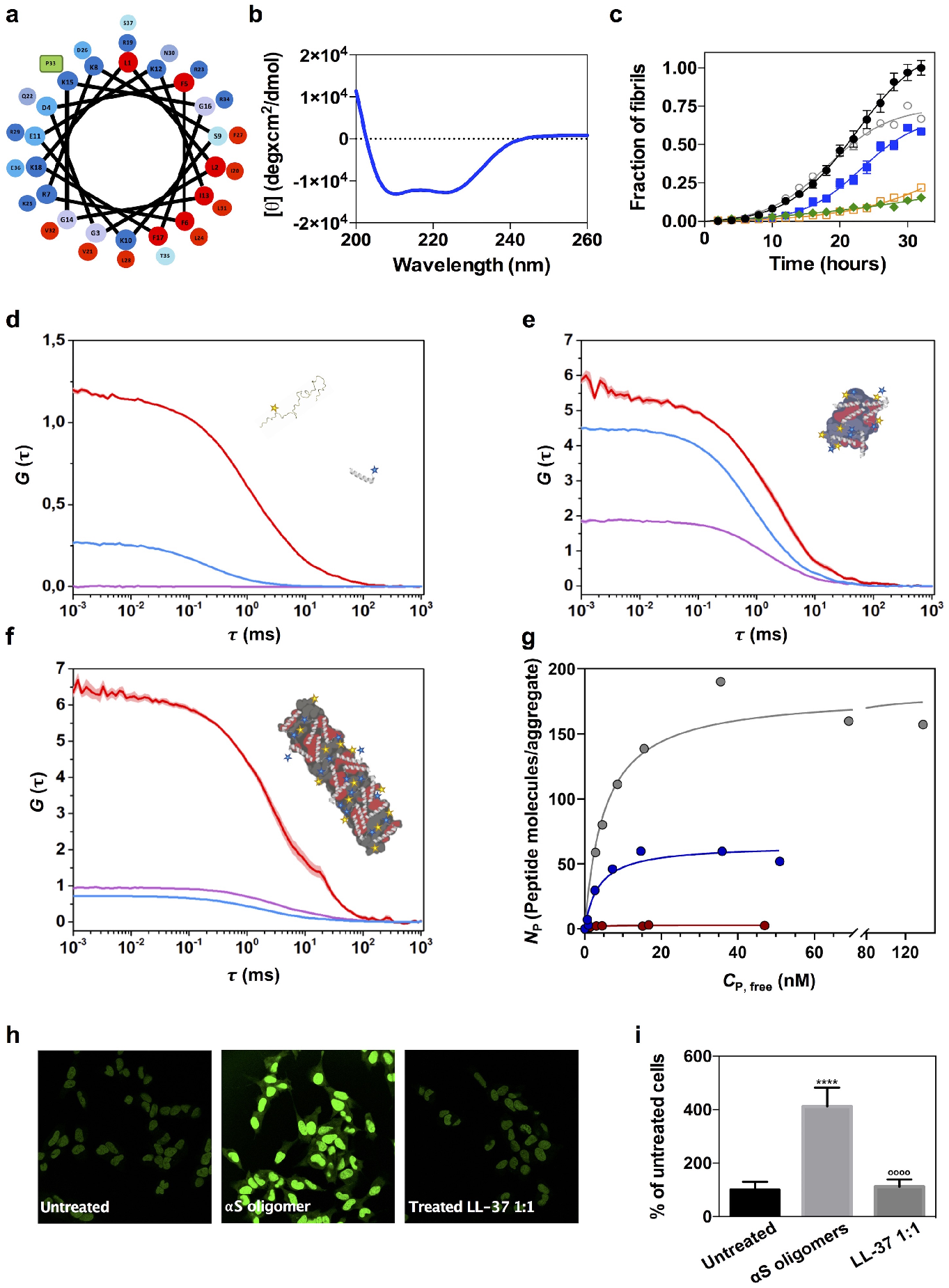
Characterization of the interaction of LL-37 with the αS pathogenic species. (**a**) Helical wheel projection of LL-37 sequence (red, hydrophobic residues; blue pallet, hydrophilic residues; green, proline). (**b**) Far-UV circular dichroism spectra of LL-37 in PBS pH 7.4. (**c**) Aggregation kinetics of 70 μM αS and titration of the inhibitory activity of LL-37 at different concentrations: 35 μM (green), 14 μM (orange), 7 μM (blue), 3.5 μM (gray) and in the absence of peptide (black). Representative auto-correlation curves for αS and LL-37 peptide and cross-correlation curves for interacting molecules are shown in blue, red and purple lines, respectively. The amplitude (*G*) error is shown as faint colored area for the corresponding correlation curves. Samples contained (**d**) ~15 nM αS monomers and ~15 nM LL-37, (**e**) 1 nM type B* oligomers and ~5 nM LL-37 or (**f**) ~5 nM sonicated fibrils and ~5 nM PSMα3. (**g**) Titration binding curves for the interaction of LL-37 with type A* oligomers (red circles), type B* oligomers (blue circles) or sonicated fibrils (grey circles) obtained by FCCS experiments, showing their corresponding analysis assuming a model of *N_P_* independent binding sites per αS aggregated species (solid lines). (**h**) Representative confocal images of SH-SY5Y cells treated with 10 μM of type B* oligomers in the presence of an equimolar concentration of LL-37. (**i**) Quantification of the intracellular ROS of the experiment displayed in panel h. Error bars represent the standard deviation (SD). **** p-value < 0.0001 relative to untreated cells. ^oooo^ p-value < 0.0001 relative to cells treated with αS type B* oligomers.

Whether LL-37 is actually involved or not in the pathogenesis of PD remains unexplored. However, our results suggest that small peptides able to interact actively with αS pathogenic species might cohabitate with this protein in tissues relevant to the disease. These human peptides would display, in principle, low immunogenicity and may open novel avenues for PD treatment, either by their direct administration or by stimulating their endogenous expression.

## Discussion

Because of its involvement in PD and other synucleinopathies, αS aggregation remains a promising target for therapeutic intervention. Herein, we propose a novel strategy for targeting the αS species behind the onset of these neurodegenerative diseases selectively. By binding to αS toxic oligomers and fibrils, the described collection of peptides inhibits the progression of αS aggregation, while suppressing the toxicity of pathogenic intermediates. Importantly, because the binding determinants are structurally encoded, these peptidic molecules do not recognize monomeric aS. From a therapeutic perspective, this is a significant advantage, as they are not expected to interfere with the physiological functions of the soluble protein. Furthermore, the avidity of these peptides for early non-toxic oligomers is more than one order of magnitude lower than the one for toxic oligomers and fibrils, indicating that they are extremely selective for the αS pathogenic species.

To the best of our knowledge, this is the first time that αS species-specific biomolecules have been rationally predicted, identified, and engineered. PSMα3 is a *first-in-class* hit molecule that sets the ground for a new generation of leads for disease modification in PD and other synucleinopathies. The requisites for a high peptide binding specificity and affinity are relatively simple: hydrophobic and positively charged surfaces with opposed orientations in the space. This is best exemplified by Scaffold_19, a short and low complexity peptide that fulfills those conditions. This defined SAR, usually absent or difficult to dissect in small molecules, should help in the development and diversification of novel candidates with increased activities employing protein engineering.

Many small bioactive peptides are derived from larger precursors and generated after proteolytic cleavage^41^. In some cases, these peptides are encrypted inside globular proteins, and their processing results in the manifestation of a new biological function. LL-37 is a cathelicidin-derived peptide constitutively expressed in the human brain^42^. Here, we show that LL-37 is a tight binder of αS pathogenic assemblies, with exceptional anti-aggregation and cytoprotective properties. LL-37 has been reported to inhibit the amyloid aggregation of two other disease-linked peptides, Aβ-42^43^ and IAPP^44^. However, the mechanism behind this activity is drastically different from the one we describe here since LL-37 binds to Aβ-42 and IAPP monomers, and the interaction is driven by a certain degree of sequence homology between short linear stretches in these molecules. Thus, it is assumed that LL-37 binds to Aβ-42 and IAPP in an, at least, partially unfolded conformation. Despite speculative, it is tempting to suggest that LL-37 is a first representative molecule of a previously unexplored regulatory system, where small peptides act as silent guardians of the proteome by targeting aggregation-prone proteins.

Apart from their therapeutic implications, the ability of the amphipathic cationic helical peptides to bind to αS toxic species with high affinity and selectivity might find a direct application in diagnosis. The presence of αS aggregates in biofluids is considered a biomarker for PD and other synucleopathies^45, 46^. However, current detection methods are not specific and sensitive enough for their clinical implementation. For instance, ELISA approaches based on the so-called conformation-specific antibodies, perform better when the detection is normalized relative to the total levels of αS or when the same epitope-blocking antibody is used for both capture and detection^47^. This indicates that a major limitation of these methods is the unwanted cross-reaction of the antibodies with the large excess of αS monomer in the fluid. In contrast, the peptides we describe here do not interact with monomeric aS, and the presence of up to 500-fold excess of monomeric αS does not interfere with the detection of nanomolar amounts of toxic oligomers. This conformational specificity, together with their close-to-antibody affinities, make of these molecules powerful tools for the implementation of diagnostic strategies in which they may act as nanosensors of pathogenic αS species in biological fluids.

Overall, the molecular entities we describe in this work represent a new conceptual framework to develop therapeutic and diagnostic strategies for the synucleinopathies.

## Acknowledgements

This work was funded by the Spanish Ministry of Economy and Competitiveness (MINECO) BIO2016-78310-R to S.V, BIO2017-91475-EXP and by ICREA, ICREA-Academia 2015 to S.V. N.C. was supported by MINECO RYC-2012-12068, MINECO/FEDER, EU BFU2015-64119-P and MICIU/FEDER, EU PGC2018-096335-B-100. J.S. was supported by the Spanish Ministry of Science and Innovation via a doctoral grant (FPU17/01157). We thank the members of the Microscopy Services of the UAB for their assistance. We also thank Dr. Evangelos Sisamakis, PicoQuant, for insightful discussion.

## Authors contribution

S.V. conceived the project. J.S., I.P. and S.V. designed all the peptide sequences. J.S., S.P-D. and J.P. performed in vitro experiments. J.S. and S.N. designed, conducted and analyzed cellular experiments. P.G. and N.C. performed and analyzed time-resolved fluorescence spectroscopy experiments. J.S., I.P. and S.V. analyzed the data. J.S., P.G., N.C., I.P. and S.V. prepared the manuscript with contributions from all the authors.

## References

1. Spillantini, M. G. et al. Alpha-synuclein in Lewy bodies. Nature 388, 839–840, (1997).

2. Winner, B. et al. In vivo demonstration that alpha-synuclein oligomers are toxic. Proc Natl Acad Sci U S A 108, 4194–4199 (2011).

3. Spillantini, M. G. & Goedert, M. The alpha-synucleinopathies: Parkinson’s disease, dementia with Lewy bodies, and multiple system atrophy. Ann N Y Acad Sci 920, 16–27 (2000).

4. McCann, H., Stevens, C. H., Cartwright, H. & Halliday, G. M. alpha-Synucleinopathy phenotypes. Parkinsonism Relat Disord 20 Suppl 1, S62–67 (2014).

5. Goedert, M., Jakes, R. & Spillantini, M. G. The Synucleinopathies: Twenty Years On. J Parkinsons Dis 7, S51–S69 (2017).

6. Kalia, L. V. & Lang, A. E. Parkinson’s disease. Lancet 386, 896–912 (2015).

7. Wong, Y. C. & Krainc, D. alpha-synuclein toxicity in neurodegeneration: mechanism and therapeutic strategies. Nat Med 23, 1–13 (2017).

8. Dehay, B. et al. Targeting alpha-synuclein for treatment of Parkinson’s disease: mechanistic and therapeutic considerations. Lancet Neurol 14, 855–866 (2015).

9. Pujols, J. et al. Small molecule inhibits alpha-synuclein aggregation, disrupts amyloid fibrils, and prevents degeneration of dopaminergic neurons. Proc Natl Acad Sci U S A 115, 10481–10486 (2018).

10. Wagner, J. et al. Anle138b: a novel oligomer modulator for disease-modifying therapy of neurodegenerative diseases such as prion and Parkinson’s disease. Acta Neuropathol 125, 795–813 (2013).

11. Kurnik, M. et al. Potent alpha-Synuclein Aggregation Inhibitors, Identified by High-Throughput Screening, Mainly Target the Monomeric State. Cell Chem Biol 25, 1389–1402 e1389 (2018).

12. Jarrett, J. T. & Lansbury, P. T., Jr. Amyloid fibril formation requires a chemically discriminating nucleation event: studies of an amyloidogenic sequence from the bacterial protein OsmB. Biochemistry 31, 12345–12352 (1992).

13. Fusco, G. et al. Structural basis of membrane disruption and cellular toxicity by alpha-synuclein oligomers. Science 358, 1440–1443 (2017).

14. Grey, M., Linse, S., Nilsson, H., Brundin, P. & Sparr, E. Membrane interaction of alpha-synuclein in different aggregation states. J Parkinsons Dis 1, 359–371 (2011).

15. Froula, J. M. et al. Defining alpha-synuclein species responsible for Parkinson’s disease phenotypes in mice. J Biol Chem 294, 10392–10406 (2019).

16. Cremades, N. et al. Direct observation of the interconversion of normal and toxic forms of alpha-synuclein. Cell 149, 1048–1059 (2012).

17. Cremades, N. & Dobson, C. M. The contribution of biophysical and structural studies of protein self-assembly to the design of therapeutic strategies for amyloid diseases. Neurobiol Dis 109, 178–190 (2018).

18. Lee, J. E. et al. Mapping Surface Hydrophobicity of alpha-Synuclein Oligomers at the Nanoscale. Nano Lett 18, 7494–7501 (2018).

19. Chen, S. W. et al. Structural characterization of toxic oligomers that are kinetically trapped during alpha-synuclein fibril formation. Proc Natl Acad Sci U S A 112, E1994–2003 (2015).

20. Chen, S. W. & Cremades, N. Preparation of alpha-Synuclein Amyloid Assemblies for Toxicity Experiments. Methods Mol Biol 1779, 45–60 (2018).

21. Cremades, N., Chen, S. W. & Dobson, C. M. Structural Characteristics of alpha-Synuclein Oligomers. Int Rev Cell Mol Biol 329, 79–143 (2017).

22. Garner, J. & Harding, M. M. Design and synthesis of alpha-helical peptides and mimetics. Org Biomol Chem 5, 3577–3585 (2007).

23. Yakimov, A. P., Afanaseva, A. S., Khodorkovskiy, M. A. & Petukhov, M. G. Design of Stable alpha-Helical Peptides and Thermostable Proteins in Biotechnology and Biomedicine. Acta Naturae 8, 70–81 (2016).

24. Marinelli, P., Pallares, I., Navarro, S. & Ventura, S. Dissecting the contribution of Staphylococcus aureus alpha-phenol-soluble modulins to biofilm amyloid structure. Sci Rep 6, 34552, (2016).

25. Munoz, V. & Serrano, L. Elucidating the folding problem of helical peptides using empirical parameters. Nat Struct Biol 1, 399–409, (1994).

26. Bacia, K. & Schwille, P. Practical guidelines for dual-color fluorescence cross-correlation spectroscopy. Nat Protoc 2, 2842–2856, (2007).

27. Schwille, P., Meyer-Almes, F. J. & Rigler, R. Dual-color fluorescence cross-correlation spectroscopy for multicomponent diffusional analysis in solution. Biophys J 72, 1878–1886 (1997).

28. Tosatto, L. et al. Single-molecule FRET studies on alpha-synuclein oligomerization of Parkinson’s disease genetically related mutants. Sci Rep 5, 16696 (2015).

29. Nath, S., Meuvis, J., Hendrix, J., Carl, S. A. & Engelborghs, Y. Early aggregation steps in alpha-synuclein as measured by FCS and FRET: evidence for a contagious conformational change. Biophys J 98, 1302–1311 (2010).

30. Whiten, D. R. et al. Single-Molecule Characterization of the Interactions between Extracellular Chaperones and Toxic alpha-Synuclein Oligomers. Cell Rep 23, 3492–3500 (2018).

31. Kruger, D., Ebenhan, J., Werner, S. & Bacia, K. Measuring Protein Binding to Lipid Vesicles by Fluorescence Cross-Correlation Spectroscopy. Biophys J 113, 1311–1320 (2017).

32. Perni, M. et al. Multistep Inhibition of alpha-Synuclein Aggregation and Toxicity in Vitro and in Vivo by Trodusquemine. ACS Chem Biol 13, 2308–2319 (2018).

33. Ludtmann, M. H. R. et al. alpha-synuclein oligomers interact with ATP synthase and open the permeability transition pore in Parkinson’s disease. Nat Commun 9, 2293 (2018).

34. Gautier, R., Douguet, D., Antonny, B. & Drin, G. HELIQUEST: a web server to screen sequences with specific alpha-helical properties. Bioinformatics 24, 2101–2102 (2008).

35. Schymkowitz, J. et al. The FoldX web server: an online force field. Nucleic Acids Res 33, W382–388 (2005).

36. Zamyatnin, A. A., Borchikov, A. S., Vladimirov, M. G. & Voronina, O. L. The EROP-Moscow oligopeptide database. Nucleic Acids Res 34, D261–266 (2006).

37. Durr, U. H., Sudheendra, U. S. & Ramamoorthy, A. LL-37, the only human member of the cathelicidin family of antimicrobial peptides. Biochim Biophys Acta 1758, 1408–1425 (2006).

38. Brundin, P. & Melki, R. Prying into the Prion Hypothesis for Parkinson’s Disease. J Neurosci 37, 9808–9818, doi:10.1523/JNEUROSCI.1788-16.2017 (2017).

39. Braak, H., Rub, U., Gai, W. P. & Del Tredici, K. Idiopathic Parkinson’s disease: possible routes by which vulnerable neuronal types may be subject to neuroinvasion by an unknown pathogen. J Neural Transm (Vienna) 110, 517–536 (2003).

40. Lionnet, A. et al. Does Parkinson’s disease start in the gut? Acta Neuropathol 135, 1–12, doi:10.1007/s00401-017-1777-8 (2018).

41. Kastin, A. J. Handbook of biologically active peptides. (Elsevier, 2013).

42. Burton, M. F. & Steel, P. G. The chemistry and biology of LL-37. Nat Prod Rep 26, 1572–1584 (2009).

43. De Lorenzi, E. et al. Evidence that the Human Innate Immune Peptide LL-37 may be a Binding Partner of Amyloid-beta and Inhibitor of Fibril Assembly. J Alzheimers Dis 59, 1213–1226 (2017).

44. Armiento, V. et al. The human cathelicidin LL-37 is a nanomolar inhibitor of amyloid self-assembly of islet amyloid polypeptide (IAPP). Angew Chem Int Ed Engl (2020).

45. Tokuda, T. et al. Detection of elevated levels of alpha-synuclein oligomers in CSF from patients with Parkinson disease. Neurology 75, 1766–1772 (2010).

46. Zhao, H. et al. AlphaLISA detection of alpha-synuclein in the cerebrospinal fluid and its potential application in Parkinson’s disease diagnosis. Protein Cell 8, 696–700 (2017).

47. Bengoa-Vergniory, N., Roberts, R. F., Wade-Martins, R. & Alegre-Abarrategui, J. Alpha-synuclein oligomers: a new hope. Acta Neuropathol 134, 819–838 (2017).

