## Supplementary information for "α-Helical peptidic scaffolds to target α-synuclein pathogenic species with high affinity and selectivity"

**a**

| Residue | Wt | 1 | 2 | 3 | 4 | 5 | 6 | 7 | 8 | 9 | 10 | 11 | 12 | 13 | 14 | 15 | 16 | 17 | 18 | 19 | 20 | 21 | 22 |
|---|---|---|---|---|---|---|---|---|---|---|---|---|---|---|---|---|---|---|---|---|---|---|---|
| α-helical predicted content | 2.6 | 3.2 | 1.8 | 1.8 | 1.9 | 1.1 | 0.9 | 0.4 | 0.6 | 0.4 | 0.4 | 0.7 | 0.7 | 0.6 | 0.6 | 0.5 | 0.8 | 1.4 | 2.0 | 2.9 | 2.3 | 2.5 | 2.9 |

**b**

**PSMα3** MEFVAKLFKFFKDLLGKFLGNN
**dPSMα3** MEFVAKLFPFPKDLLGKFLGNN
* *

**c** **d**

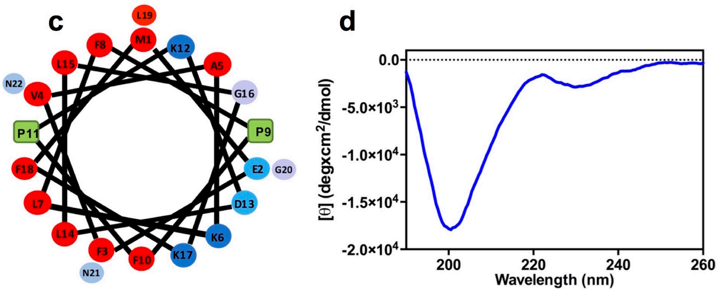


**Supplementary Figure 1. Design of a non-amphipathic PSMα3 variant (dPSMα3).** (**a**) Computational proline scanning. Predicted α-helical propensity according to the AGADIR score; higher values indicate higher predicted α-helical propensity. (**b**) Sequence alignment of PSMα3 and dPSMα3. (**c**) Helical wheel projection of dPSMα3 sequences showing the theorical location of the introduced prolines (green) (red, hydrophobic residues; blue pallet, hydrophilic residues depending on their character). (**d**) Far-UV circular dichroism spectrum of dPSMα3.

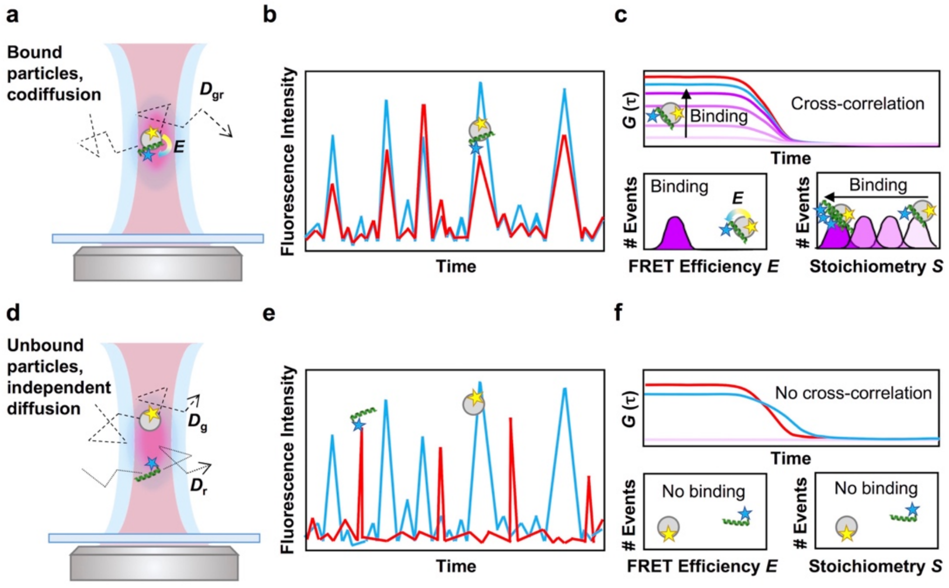


**Supplementary Figure 2. Schematic figure showing the two-color time-resolved fluorescence spectroscopy approach to characterize the binding of the peptides to the different $\alpha$S species**. The upper panels (**a-c**) illustrate a scenario where binding occurs whereas the lower panels (**d-f**) serve as an example of a non-binding scenario. (**a**) two interacting molecules labeled with a green and a red dye (depicted as a yellow or blue star, respectively) freely co-diffuse through the dual-laser confocal volume. The co-diffusion of the molecules is indicated as $D_{gr}$ while FRET between the dyes in the complex is indicated as *E*. (**b**) Illustration of a fluorescence time-trace of the co-diffusing molecules where intensity bursts of the green and red detection channels (blue and red traces, respectively) coincide in time. (**c**) the upper panel illustrates a positive cross-correlation scenario (purple lines) where the cross-correlation amplitude (*G*) is directly proportional to the degree of binding. The bottom left panel depicts a FRET efficiency (*E*) distribution from interacting molecules while the bottom right panel shows the green dye-to-red dye (donor-to-acceptor) stoichiometry of those interacting particles and shows how the stoichiometry decreases with an increasing binding degree. (**d-f**) illustrate the same parameters in the case where no interaction is observed.

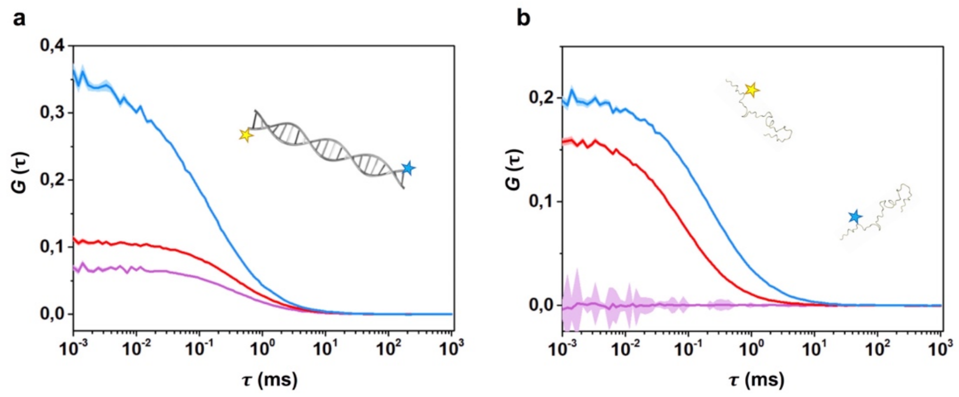


**Supplementary Figure 3. Fluorescence cross-correlation spectroscopy positive and negative control.** Auto-correlation curves of AF488 (blue) and Atto647N (red) and cross-correlation curves (purple) of samples containing (**a**) 10 nM of doubly-labelled dsDNA molecule or (**b**) 15 nM of non-interacting AF488-$\alpha$S and Atto647N-$\alpha$S (15 nM each). The amplitude (*G*) error is shown as faint colored area for the corresponding correlation curves.

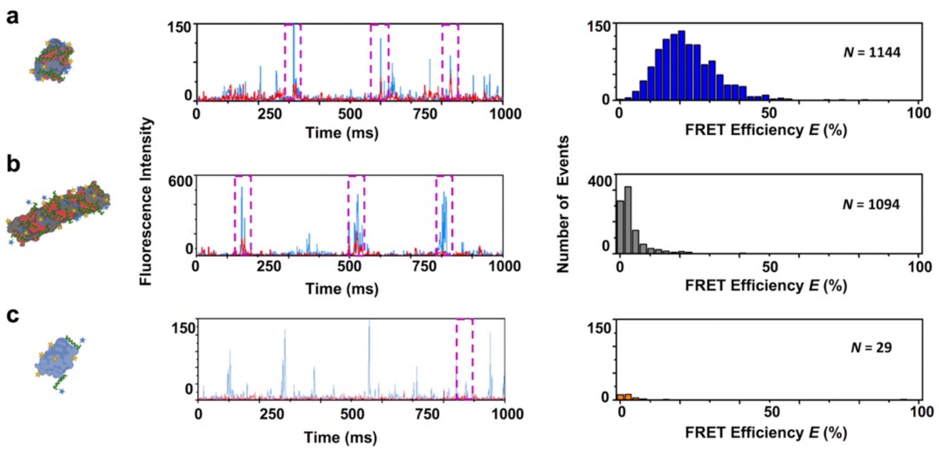


**Supplementary Figure 4. $\alpha$S-PSM$\alpha$3 binding analyzed by Fluorescent single-particle spectroscopy.** Representative intensity time traces (left panels) and intensity-calculated FRET efficiency histograms (right panels) for samples containing (**a**) ~1 nM $\alpha$S type B* oligomers and ~5 nM PSM$\alpha$3, (**b**) ~5 nM $\alpha$S fibrils and ~5 nM PSM$\alpha$3 and (**c**) 1 nM $\alpha$S type A* oligomers and ~5 nM PSM$\alpha$3. In the intensity traces, events displaying both donor and acceptor intensities above $\alpha$S monomer threshold (see materials and methods) are shown in purple dashed boxes. These events were then used to calculate the intensity-based FRET efficiency *E* histograms. The total number of FRET events, *N*, used to calculate each histogram is shown in each panel. These results show, directly from the intensity raw data, the high avidity of both PSM$\alpha$3 for either type B* oligomers or fibrils (**a-b**) and the low ability to bind non-toxic aggregated species like the type A* oligomers (c).

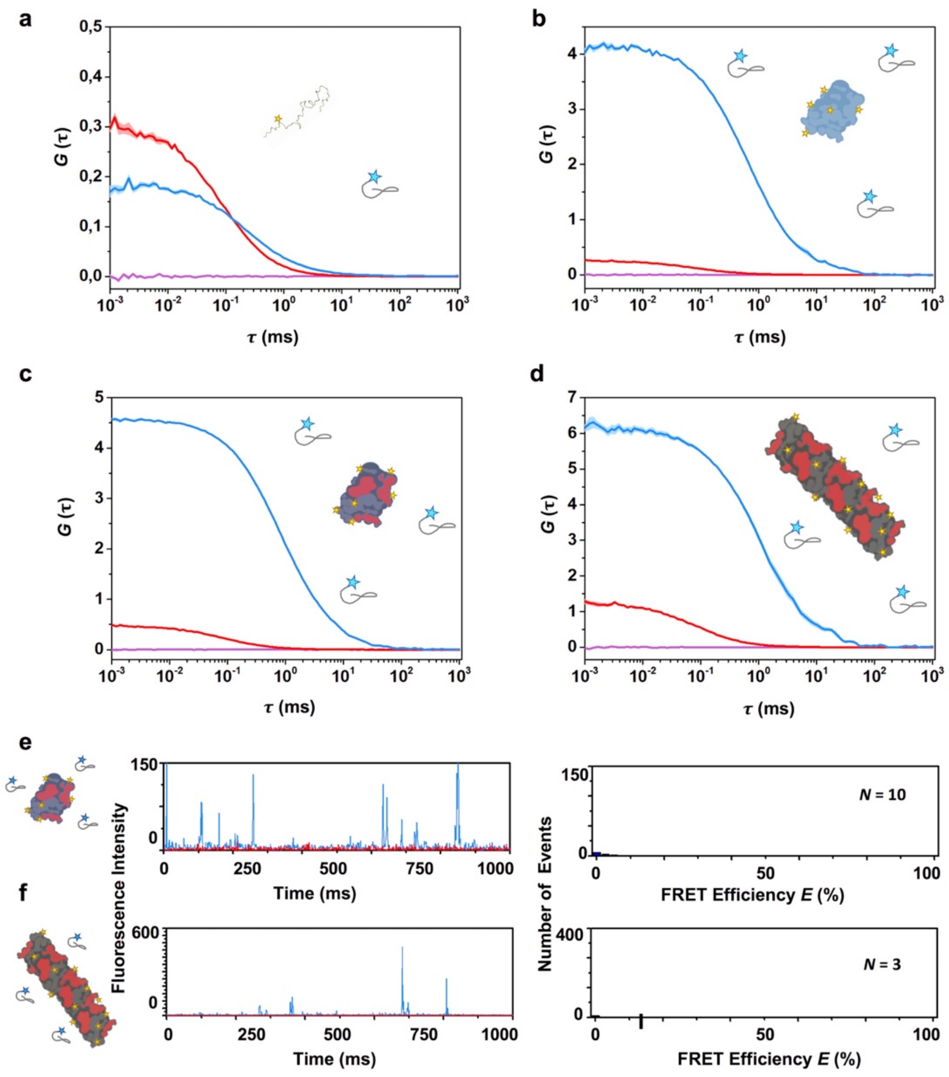


**Supplementary Figure 5. Interaction of dPSM$\alpha$3 with the different $\alpha$S species.** (**a-d**) Auto-correlation curves for $\alpha$S and dPSM$\alpha$3 and cross-correlation curves for interacting molecules are shown in blue, red and purple lines, respectively. The amplitude ($G$) error is shown in faint blue, red and purple, respectively. ~15 nM $\alpha$S monomer (**a**), ~1 nM type A* (**b**), type B* (**c**) oligomers and sonicated fibrils (**d**) were allowed to interact with ~15 nM dPSM$\alpha$3. No cross-correlation is observed in any case. (**e-f**) $\alpha$S-dPSM$\alpha$3 binding analyzed by single-particle fluorescent spectroscopy. Representative intensity time traces (left panels) and intensity-calculated FRET efficiency histograms (right panels) for samples containing (**e**) ~1 nM $\alpha$S type B* oligomers and ~5 nM dPSM$\alpha$3, (**f**) ~5 nM $\alpha$S fibrils and ~5 nM dPSM$\alpha$3. FRET single-particle fluorescence data directly show the inability of dPSM$\alpha$3 to interact with either type B* oligomers or fibrils (**e, f**), as just few, if any, events were observed in comparison to the experiments with the PSM$\alpha$3 peptide.

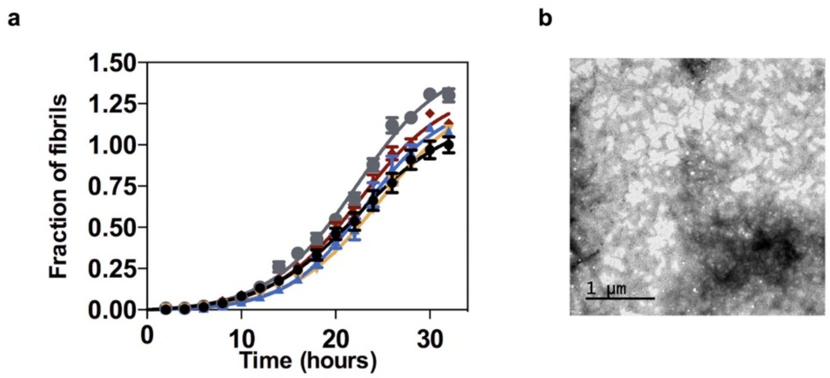


**Figure S6. Effect of dPSMα3 on *in vitro* αS amyloid fibrillation.** (**a**) Aggregation kinetics of 70 µM αS and titration of the inhibitory activity of dPSMα3 at different concentrations: 35 µM (green), 14 µM (orange), 7 µM (blue), 3.5 µM (gray) and in the absence of dPSMα3 (black). Error bars represent the standard error of the mean (SEM). (**b**) TEM micrograph of the end point of the aggregation kinetics in the presence of 70 µM of dPSMα3.

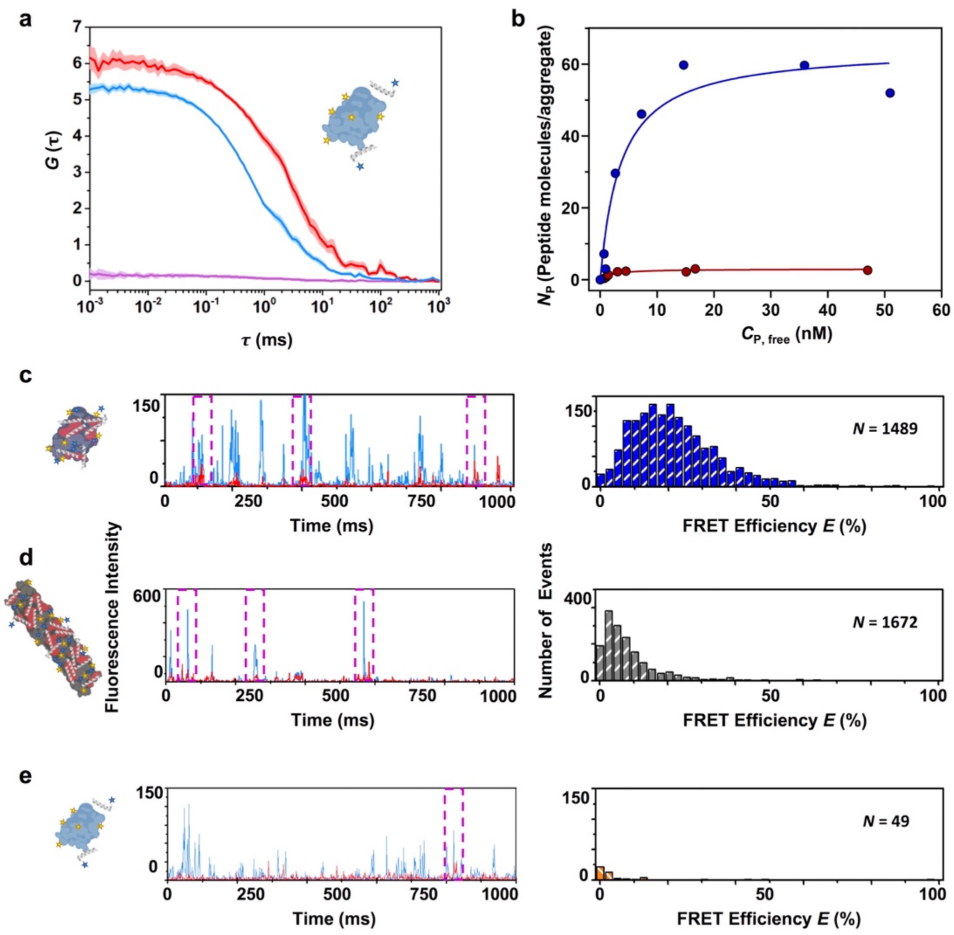


**Supplementary Figure 7. Interaction of LL-37 with αS aggregates by FCCS and fluorescent single-particle spectroscopy.** (**a**) Auto-correlation curves for αS (blue) and LL-37 (red) and cross-correlation curve for the interacting molecules (purple) in samples containing ~1 nM type-A* oligomers and ~5 nM LL-37 peptide. The amplitude (*G*) error is shown as faint colored area for the corresponding correlation curves. (**b**) Titration binding curves for the interaction of LL-37 with type-A* oligomers (red circles) or type-B* oligomers (blue circles) obtained by FCCS experiments, showing their corresponding analysis assuming a model of $N_P$ independent binding sites per αS aggregated species (solid lines). (**c-e**) αS-LL-37 binding analyzed by Fluorescent single-particle spectroscopy. Representative intensity time traces (left panels) and intensity-calculated FRET efficiency histograms (right panels) for samples containing (**c**) ~1 nM αS type-B* oligomers and ~5 nM LL-37, (**d**) ~5 nM αS fibrils and ~5 nM LL-37 and (**e**) ~1 nM αS type-A* oligomers and ~5 nM LL-37.

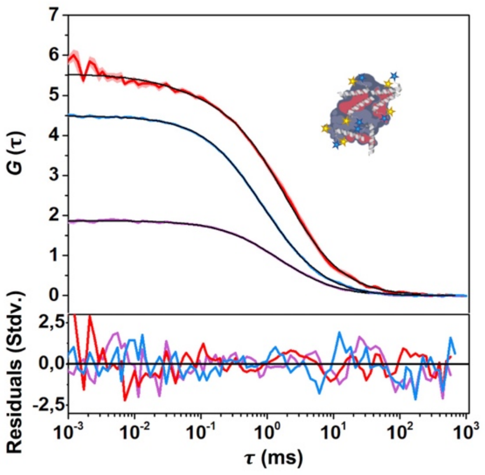


**Supplementary Figure 8. Fitting of correlation and cross-correlation functions.** Representative auto-correlation and cross-correlation curves of a sample of 1 nM type-B* oligomers and 5 nM PSM$\alpha$3 peptide are shown in blue, red and purple lines, respectively. The amplitude (*G*) error is shown as faint colored area for the corresponding correlation curves. Best fits to 1-diffusion component (cross-correlation) or 2-diffusion component (auto-correlations) simple diffusion models are shown as black lines. The residual analysis of the best fits is also shown as standard deviation in colored lines for each correlation curve fit.

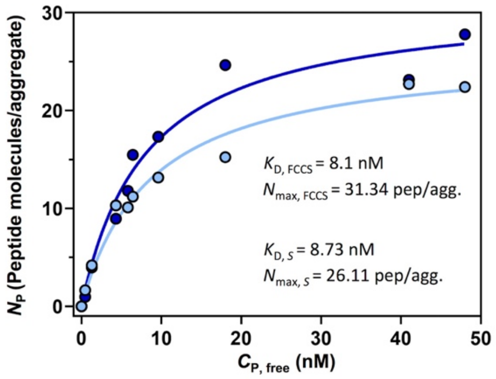


**Supplementary Figure 9. Comparison of the titration binding curves of αS type B* oligomers with PSMα3 peptide obtained by FCCS or two-color coincidence single-particle fluorescence analysis.** The number of peptide molecules bound to one oligomer ($N_P$) at increasing peptide concentrations was calculated independently by FCCS (dark blue circles) or stoichiometry analysis in two color coincidence single-particle fluorescence experiments (light blue circles) yielding very similar titration binding curves that resulted in very similar binding parameters when analyzed using a model of *n* identical and independent binding sites per αS aggregated species (solid lines). The fitted parameters $K_D$ and $N_{max}$ are also shown for each analytical approach. These results show how two different analytical methods, one which correlates whole time traces and other one which analyses single fluorescent bursts, can be applied to obtain very similar binding parameters, thus validating our strategy.

**Supplementary Table 1. Computational analysis of redesigned variants.**

| Name | Mutacion | Sequence | Activity | *H** | $\mu_H$** | AGADIR | Net charge | Change in FoldX stability (kcal/mol) |
|---|---|---|---|---|---|---|---|---|
| **PSMα3** | - | MEFVAKLFKFFKDLLGKFLGNN | +++ | 0.54 | 0.56 | 2.65 | 2 | - |
| **dPSMα3** | K9P_F11P | MEFVAKLFPFPKDLLGKFLGNN | - | 0.57 | 0.44 | 0.40 | 1 | 3.67 |
| **All_Leu** | Hydrophobic face to Leu | LELLAKLLKLLKDLLGKLLGNN | +++ | 0.57 | 0.58 | 66.14 | 2 | -1.46 |
| **All_Leu19** | Hydrophobic face to Leu without 3 C-ter residues | LELLAKLLKLLKDLLGKLL | +++ | 0.72 | 0.70 | 65.17 | 2 | -1.46 |
| **Scaffold_19** | Hydrophobic face to Leu without 3 C-ter residues A5E_G16K | LELLEKLLKLLKDLLKKLL | +++ | 0.62 | 0.77 | 77.68 | 2 | -1.03 |
| **Anionic scaffold** | Scaffold19 K6E_K12E | LELLEELLKLLEDLLKKLL | - | 0.65 | 0.75 | 78.26 | -2 | 0.65 |

* *H* indicates the mean hydrophobicity of the peptides.
** $\mu_H$ indicates the helical hydrophobic moment of the peptides.

**Supplementary Table 2. Identified human peptide candidates.** The screening of the human peptides database (EROP-Moscow) for cationic peptides with more than 10 residues, an AGADIR value > 2 and a helical hydrophobic moment ($\mu_H$) > 0.2.

| Peptide sequence | AGADIR | $\mu_H$ | Cysteines |
|---|---|---|---|
| >E02311 \| ANTIMICROBIAL PEPTIDE CATHELICIDIN LL37 \| HUMAN (HOMO SAPIENS), COMMOM CHIMPANZEE (PAN TROGLODYTES)<br><br>LLGDFFRKSKEKIGKEFKRIVQRIKDFLRNLVPRTES | 5.10 | 0.521 | No |
| >E02310 \| ANTIMICROBIAL PEPTIDE CATHELICIDIN FALL 39 \| HUMAN (HOMO SAPIENS), COMMOM CHIMPANZEE (PAN TROGLODYTES)<br><br>FALLGDFFRKSKEKIGKEFKRIVQRIKDFLRNLVPRTES | 4.92 | 0.529 | No |
| >E19967 \| SALUSIN BETA \| HUMAN (HOMO SAPIENS)<br><br>AIFIFIRWLLKLGHHGRAPP | 2.14 | 0.306 | No |
| >E06260 \| ANAPHYLATOXIN C3A PEPTIDE LGE27 \| HUMAN (HOMO SAPIENS)<br><br>LGEACKKVFLDCCNYITKLRRQHARAS | 5.63 | 0.493 | Yes |
| >E06257 \| ANAPHYLATOXIN C3A PEPTIDE SLG25 \| HUMAN (HOMO SAPIENS)<br><br>SLGEACKKVFLDCCNYITELRRQHA | 4.72 | 0.48 | Yes |
| >E01232 \| MELANIN CONCENTRATING HORMONE \| RAT (RATTUS NORVEGICUS), HUMAN (HOMO SAPIENS), MOUSE (MUS MUSCULUS)<br><br>DFDMLRCMLGRVYRPCWQV | 3.82 | 0.406 | Yes |
| >E06261 \| ANAPHYLATOXIN C3A PEPTIDE CNY21 \| HUMAN (HOMO SAPIENS)<br><br>CNYITELRRQHARASHLGLAR | 4.17 | 0.241 | Yes |
| >E05394 \| BETA DEFENSIN 4 \| HUMAN (HOMO SAPIENS)<br><br>EFELDRICGYGTARCRKKCRSQEYRIGRCPNTYACCLRKWDESLLNRTKP | 3.09 | 0.32 | Yes |
| >E04240 \| BETA DEFENSIN 6, HBD6 \| HUMAN (HOMO SAPIENS)<br><br>FFDEKCNKLKGTCKNNCGKNEELIALCQKSLKCCRTIQPCGSIID | 3.23 | 0.231 | Yes |
