## Supplementary material for "α-Helical peptidic scaffolds to target α-synuclein pathogenic species with high affinity and selectivity": Methods

### αS expression and purification

Human αS was expressed and purified as previously described[1,2]. *Escherichia coli* BL21 (DE3) cells containing a pET21a plasmid encoding the αS gene were grown in LB medium supplemented with 100 µM/mL ampicillin. Protein expression was induced at an optical density of 0.8 (600 nm) 1 mM isopropyl β-D-thiogalactopyranoside (IPTG) for 4 h. Cells were harvested by centrifugation and washed up by resuspension and centrifugation in PBS pH 7.4. Next, pellets were resuspended in 50 mL per culture liter in lysis buffer (50 mM Tris pH 8, 150 mM NaCl, 1µg/mL pepstatin, 20 µg/mL aprotinin, 1 mM benzamidine, 1 mM PMSF, 1 mM EDTA and 0.25 mg/mL lysozyme) and sonicated using a LabSonic®U sonicator (B. Braun Biotech International, Melsungen, Germany). Samples were boiled during 10 minutes at 95 °C and centrifugated at 20,000g at 4 °C for 40 minutes. The soluble fraction was treated with 136 µL/mL of 10 % w/v streptomycin sulfate and 228 µL/mL of pure acetic acid. Upon centrifugation, soluble extracts were fractionated by adding 1:1 of saturated ammonium sulfate and resuspending the insoluble fraction with 50 % ammonium sulfate. The pellet was resuspended in 100 mM pH 8 ammonium acetate (5 mL per culture liter) and pure EtOH 1:1 (v/v) and harvested by centrifugation. The insoluble fraction was resuspended in Tris 20 mM pH 8, filtered with a 0.22 µm filter and loaded into an anion exchange column HiTrap Q HP (GE Healthcare, Chicago, USA) coupled to an ÄKTA purifier high performance liquid chromatography system (GE Healthcare, Chicago, USA). Tris 20 mM pH 8 and Tris 20 mM pH 8, NaCl 1 M were used as buffer A and buffer B. αS was eluted using a using a step gradient: Step 1: 0 %–20 % buffer B, 5 cv; Step 2: 20 %–45 % buffer B, 11 cv; Step 3: 100 % buffer B, 5 cv. Purified αS was dialyzed against 5L ammonium acetate 50 mM in two steps; 4 h and overnight. Finally, protein purity was addressed using 15 % SDS-PAGE. The purest fractions were lyophilized and stored at -80 °C. For the experiments, αS lyophilized aliquots were resuspended to a final concentration of 210 μM using PBS pH 7.4 and filtered using 0.22 μm filters. αS concentration was determined measuring the absorbance at 280 nm and using the extinction coefficient 5,960 $M^{-1}$ $cm^{-1}$.

**Peptide preparation**

PSMα3, dPSMα3, All_Leu, All_Leu_19, Scaffold_19, Anionic Scaffold and LL-37 were purchased from Synpeptide (Shanghai, China) with a purity >95 %. Single cysteine containing variants were purchased from Genscript (Piscataway, USA) with a purity >95 %. LL-37 was diluted in Milli-Q sterilized water, divided into aliquots and lyophilized. Cysteine containing peptides were resuspended in PBS pH 7.4, 5 mM TCEP and subsequently labeled with the corresponding fluorophore. PSMα3, dPSMα3, All_Leu, All_Leu_19, Scaffold_19 and Anionic Scaffold were dissolved in a 1:1 mixture of trifluoroacetic acid and hexafluoroisopropanol and sonicated for 10 minutes. Stock solutions were divided into aliquots and vacuum dried with a SpeedVac (Thermo Fisher Scientific, Waltham, USA) and stored at -80 °C until assayed. Peptide aliquots were resuspended in pure Milli-Q water prior their use.

**αS and peptide labeling**

Site-specific labeling of αS was performed in an αS variant with a single engineered cysteine at position 122 (αS N122C). This variant was expressed and purified as previously described[3,4]. The protein was labeled with maleimide-modified Alexa Fluor 488 (AF488) (Invitrogen, Carlsbad, USA) for 15-20 h at 4 °C in the dark. After quenching the reaction with 10 mM DTT, free unreacted dye in the protein solution was subsequently separated using a P10 desalting column (GE Healthcare, Waukesha, USA), and the labeled protein solution was flash frozen with liquid nitrogen and stored at -80 °C. The different peptides, PSMα3, dPSMα3 and LL-37, were labeled at a single engineered cysteine at the N-terminus with maleimide-modified Atto647N (ATTO-TEC, Siegen, Germany). The same labeling and purification strategy were followed as for αS, although in this case the unreacted free dye was removed from the protein solution using a polyacrylamide desalting column (Thermo Fisher Scientific, Waltham, USA). Two cleaning steps were required to remove completely the free dye from the labeled peptide solution.

**Preparation of the different isolated αS aggregates samples**

Oligomeric samples were prepared as previously described[5,6]. For the isolation of type B* oligomers purified αS was dialyzed against Milli-Q water and lyophilized for 48 h in aliquots of 6 mg. The aliquots were resuspended in 500 µL of PBS pH 7.4 to a final concentration of ca. 800 µM, filtered through 0.22 µm filters and incubated at 37 °C without agitation for 20-24 h. The sample was then ultracentrifuged at 288,000g in a SW55Ti Beckman rotor, in order to remove any possible fibrillar species formed during the incubation, and later filtered by four consecutive cycles of filtration through 100 kDa centrifuge filters (Merck, Darmstadt, Germany) in order to remove the great excess of monomeric protein from the oligomeric solution. Type A* oligomers were generated by incubating 210 µM of αS in PBS pH 7.4 with ten molar equivalents of (-)-epigallocatechin-3- gallate (EGCG) (Merck, Darmstadt, Germany) for 48 h at 37 °C. The excess of compound and unreacted monomeric protein were then removed by six consecutive cycles of filtration through 100 kDa centrifuge filters (Merck, Darmstadt, Germany). The concentration of the final oligomeric solutions was determined measuring the absorbance at 280 nm and using an extinction coefficient of 5,960 $M^{-1}$ $cm^{-1}$ or absorbance at 495 nM and an extinction coefficient of 72,000 $M^{-1}$ $cm^{-1}$ for AF488-labeled oligomers. In all cases, the oligomers were kept at room temperature and were used within 3 days after their production. The fibrillar samples were produced as explained in the aggregation kinetics methodology section. The non-reacted protein and small non-fibrillar species that could be formed during the aggregation reaction were removed from the sample by 3 consecutive steps of centrifugation and resuspension of the precipitated fraction in PBS buffer at pH 7.4. Fibrils were then sonicated (1 min, 50 % cycles, 80 % amplitude in a Vibra-Cell VC130 Ultrasonic Processor (Sonics, Newton, USA) to generate fibrillar samples with a relatively homogeneous size distribution of small fibrils. The concentration of the AF488-labeled fibrillar samples was determined by subtracting the absorbance of the monomer after centrifugation at 495 nm using an extinction coefficient of 72,000 $M^{-1}$ $cm^{-1}$, with respect of the total soluble at time 0.

**Far circular dichroism analysis**

Far-UV CD spectra of the different peptide solutions were recorded on a Jasco J-815 CD spectrometer (Halifax, Canada) at 25 °C using samples of 15 μM peptide final concentration in Milli-Q water. CD signal was measured from 260 nm to 190 nm at 0.2 nm intervals, 1 nm bandwidth, 1 second of response time and a scan speed of 100 nm/min on a 0.1 cm quartz cell. 10 accumulations were recorded and averaged for each measurement. For LL-37 peptide samples, CD spectra were recorded in PBS pH 7.4, because of structural differences of this peptide in water and saline solvents.

**Time-Resolved Fluorescence Spectroscopy**

Dual-Color Time-Resolved Fluorescence Spectroscopy experiments were performed on a commercial MT200 (PicoQuant, Berlin, Germany) time-resolved fluorescence confocal microscope with a Time-Correlated Single Photon Counting (TCSPC) unit. Laser diode heads were used in Pulsed Interleaved Excitation (PIE), coupled through a single-mode waveguide and adjusted to laser powers of 6 μW (481 nm) and 5 μW (637 nm) measured after the dichroic mirror for optimal count rates while avoiding photobleaching and saturation. The coverslip was placed directly on the immersion water on top of a Super Apochromat 60x NA 1.2 objective with a correction collar (Olympus Life Sciences, Waltham, USA). A dichroic mirror of 488/640 nm (Semrock, Lake Forest, IL, USA) was chosen as the main beam splitter. Out of focus emission light was blocked by a 50 μm pinhole and the in-focus emission light was then split by a 50/50 beamsplitter into 2 detection paths. Bandpass emission filters (Semrock, Lake Forest, IL, USA) of 520/35 for the green dye (AF488) and 690/70 for the red dye (Atto647N) were used before the detectors. Single Photon Avalanche Diodes (SPADs) (Micro Photon Devices, Bolzano, Italy) served as detectors. Each measurement had an acquisition time of 1 to 3 minutes.

For FCS experiments, the effective focal volume of the green channel and its structural parameters in our system were determined using a 1 nM solution of Atto488 (ATTO-TEC GmbH, Siegen, Germany) yielding $V_{eff,\,g} = 0.51$ fL and $\kappa_g = 3.97$. Positive and negative cross-correlation controls were performed with a dual-labeled dsDNA (10 nM) and an equimolar mixture (15 nM each) of AF488- and Atto647N-labelled monomeric αS (**Supplementary Figure 3**). The positive control was also used for the determination of the red and dual-color effective focal volume and their structural parameter, yielding $V_{eff,r} = 0.1$ fL, $V_{eff,gr} = 0.091$ fL, $\kappa_r = 2.78$ and $\kappa_{gr} = 2.67$, respectively.

The AF488-labeled αS samples were diluted in PBS pH 7.4 to a final protein concentration of~5 nM in a 50 µL droplet and spotted directly onto a cover glass (Corning, Corning, USA) previously coated with a 1 mg/mL BSA solution. Atto 647N-labeled peptides were titrated into the droplet and the peptide concentration was measured individually for each experiment by autocorrelation analysis of the red dye. No significant changes in correlation amplitudes were observed over time after equilibrating the samples for 2 minutes. Experiments were performed at 20 °C and samples were covered to avoid evaporation.

Both data acquisition and analysis were performed on the commercially available software SymphoTime64 (PicoQuant, Berlin, Germany). For the oligomeric and fibrillar samples, a lower intensity threshold of 27 photons in the green dye auto correlation analysis was applied to filter out the low intensity signal arising from the monomeric αS events generated upon dilution-induced disaggregation of the aggregated samples. This threshold was calculated as 3 times the mean intensity of monomeric αS obtained from the analysis of a sample of pure αS monomers. Additionally, an upper intensity threshold was applied to auto-correlation and cross-correlation analysis to filter out any possible artifacts such as dust particles or aggregate clusters (anyway these events were very scarce): 500 photons for monomer, type A* and type B* oligomers and 1500 photons for sonicated fibrils. The PIE excitation scheme together with the TSCPC acquisition enabled the application of a lifetime-weighted filter which aided removal of background and spectral cross-talk. The corrected auto-correlations of the green and the red channel ($G_i$) were given by

$$G_{\mathrm{i}}\,(\tau) = \frac{\langle F_{\mathrm{i}}(t)\cdot F_{\mathrm{i}}\,(t+\tau)\rangle}{{F_{\mathrm{i}}}^2} - 1 \qquad \text{(Eq. 1)}$$

where $F_i\,(t)$ denotes the fluorescence intensity either the green or the red channel, $\tau$ is the correlation time and the angled brackets indicate a time average over the acquisition time. The cross-correlation ($G_x$) between the green and the red channel was given by

$$G_{\mathrm{x}}\,(\tau) = \frac{\langle F_{\mathrm{g}}(t)\cdot F_{\mathrm{r}}\,(t+\tau)\rangle}{\langle F_{\mathrm{g}}\rangle\langle F_{\mathrm{r}}\rangle} - 1 \qquad \text{(Eq. 2)}$$

Auto-correlation curves for both the green and red channel were fitted with a 2 diffusion-component model accounting for residual monomeric αS and bound and unbound peptide, respectively, using the following equation:

$$G_{\mathrm{i}}\,(\tau) = G_{\mathrm{i}}^{0}\,\frac{f_{\mathrm{i},1}}{\left(1+\frac{\tau}{\tau_{\mathrm{D_{i,1}}}}\right)\sqrt{1+\frac{\tau}{\kappa^2\ \mathrm{x}\ \tau_{\mathrm{D_{i,1}}}}}} + \frac{f_{\mathrm{i},2}}{\left(1+\frac{\tau}{\tau_{\mathrm{D_{i,2}}}}\right)\sqrt{1+\frac{\tau}{\kappa^2\ \mathrm{x}\ \tau_{\mathrm{D_{i,2}}}}}}, \qquad \text{(Eq. 3)}$$

where $G_i^0$ is the correlation amplitude at correlation time 0, $f_{i,1}$ and $f_{i,2}$ denote the fractional amplitudes of the monomeric and aggregated αS for the green channel (where i = g) and the bound and unbound peptide for the red channel (where i = r) and $\kappa^2$ is the structure parameter of the focal volume. The same applies for the diffusion terms $\tau_{D_{i,1}}$ and $\tau_{D_{i,2}}$. No correlated blinking is expected when multiple dyes are present on one particle as it is our case and therefore a blinking term was not included.

Cross-correlation amplitudes were fitted with a 1-component simple diffusion model since only one diffusion coefficient is expected for the interacting species (**Supplementary Figure 8**) using the following equation:

$$G_x(\tau) = G_x^0 \frac{1}{\left(1+\frac{\tau}{\tau_{D,x}}\right)\sqrt{1+\frac{\tau}{\kappa^2 \times \tau_{D,x}}}}. \quad \text{(Eq. 4)}$$

With the corrected green dye autocorrelation function and the mean intensity of monomeric αS, the average aggregate particle number ($N_{Ag}$) for each αS aggregated sample was estimated as $N_{Ag} = \frac{1}{G_g^0}$. The peptide concentration was calculated as $C_P = \frac{N_r}{V_{eff,r} \times N_A}$, where $N_r$ is the average number of particles in the red confocal volume, $V_{eff,r}$ is the red focal volume and $N_A$ is the Avogadro number. The cross-correlation amplitudes, $N_{Ag}$, dual-laser focal volume, $V_{eff,x}$, and peptide concentrations, $C_P$, were used for calculating the number of peptides bound to each αS species ($N_P$) and the free peptide concentration ($C_{P,Free}$) as described by Kruger and coworkers[7].

For single-burst FRET and stoichiometry analysis, an acceptor (red dye) direct excitation lower threshold based on the mean intensity of the time trace ($I_{A,mean}$+ 2×σ) was used to filter out those events without an active acceptor molecule. To further select those events arising from αS aggregates, a burst selection intensity threshold of 100 photons was used. In the FRET analysis, experimentally determined correction factors were applied: spectral cross-talk $\alpha$ was 0.004, direct excitation $\beta$ was 0.0305 and detection efficiency $\gamma$ was 0.517. Burst-wise FRET efficiency and stoichiometry were calculated as given by

$$E = \frac{F_{\mathrm{A,IE}}}{F_{\mathrm{D}} + F_{\mathrm{A,SE}}} \qquad \text{(Eq. 5)}$$

$$S = \frac{F_{\mathrm{D}} + F_{\mathrm{A,IE}}}{F_{\mathrm{D}} + F_{\mathrm{A,IE}} + F_{\mathrm{A,DE}}} \qquad \text{(Eq. 6)}$$

where $F_{\mathrm{D}}$ is the fluorescence intensity in the donor (green) channel, $F_{\mathrm{A,IE}}$ is the fluorescence intensity in the acceptor (red) channel through indirect excitation and $F_{\mathrm{A,DE}}$ is the fluorescence intensity in the acceptor (red) channel after direct excitation by PIE pulse.

Stoichiometry values were corrected for the difference in mean intensity between the monomeric αS and peptide bursts, obtained from monomeric αS-only and peptide-only measurements; the obtained mean intensity ratio $I_{\mathrm{mean},\alpha\mathrm{S}}$:$I_{\mathrm{mean,peptide}}$ was found to be 0.77. Stoichiometry distributions were fitted to a log-normal distribution to obtain the mean stoichiometry value for each measurement. The number of bound peptides per aggregate ($N_{\mathrm{P}}$) was then estimated by multiplying the mean stoichiometry value previously obtained by the mean number of αS monomers present on each aggregate as calculated empirically from the molecular brightness in FCCS experiments. The free peptide concentration ($C_{\mathrm{P,\,Free}}$) and $N_{\mathrm{P}}$ obtained by either FCCS or single-burst stoichiometry analysis were used for calculating the binding curves as described by Kruger and coworkers[7]. To obtain the dissociation constant $K_{\mathrm{D}}$

and the maximum specific binding sites $N_{max}$, the resulting binding curves were fitted to the following specific binding model with $n$ identical and independent binding sites:

$$Y = \frac{N_{\max} \cdot X}{(K_D + X)} \qquad \text{(Eq. 7)}$$

The binding curves and binding parameters obtained from either FCCS or single-burst stoichiometry analysis were compared (**Supplementary Figure 9**) and found to be remarkably similar, which validates the analysis.

**Aggregation kinetics**

αS amyloid aggregation was monitored in a 96 wells plate (non-treated) (Sarstedt, Germany) containing Teflon polyballs (1/8″ diameter) (Polysciences Europe GmbH, Eppelheim, Germany) as described by Pujols and coworkers[1]. Each well contained 150 µL solutions of 70 µM αS in PBS buffer with 40 µM thioflavin-T and the corresponding concentration of peptide. Plates were incubated at 37 °C, 100 rpm in an orbital culture shaker Max-Q 4000 (Thermo Fisher Scientific, Waltham, USA). Aggregation was analyzed every 2 h using a Victor3.0 Multilabel Reader (PerkinElmer, Waltham, USA). End-point measurements were performed after 32 h of incubation. Fluorescence intensity was measured in triplicate by exciting with a 430–450 nm filter and collecting the emission with a 480–510 nm filter. The resulting kinetics were normalized to the maximum fluorescence of the αS control (untreated).

**Transmission electron microscopy**

For electron microscopy analyses, end-point aggregated samples were sonicated for 5 min at minimum intensity in an ultrasonic bath (VWR ultrasonic cleaner) and placed onto carbon-coated copper grids and allowed to absorb for 5 min. The grids were then washed with distilled water and negative stained with 2 % (w/v) uranyl acetate for 1 minute. Finally, the excess of uranyl acetate was absorbed using ashless filter paper and the grids were left to air-dry for 15 minutes. A TEM JEM-1400 (JEOL, Peabody, USA) microscope was used operating at an accelerating voltage of 120 kV. The more representative images of each grid were selected.

**Neuroblastoma culture**

Human SH-SY5Y neuroblastoma cells (ATCC) were cultured in DMEM/F12 medium supplemented with 15 % FBS and 1xNEAA. Cells were grown at 37 °C in a 5 % $CO_2$ humidified atmosphere until an 80 % confluence for a maximum of 20 passages.

**Analysis of intracellular ROS.**

SH-SY5Y cells were seeded onto glass coverslips (Ibidi, Gräfelfing, Germany) at $0.5x10^6$ cells/mL and treated for 15 minutes with 10 μM of type B* oligomers or type B* pretreated for 15 minutes with the tested peptide (PSMα3, dPSMα3 and LL-37). Then, CellROX® Green (Invitrogen, Carlsbad, USA) at a final concentration of 5 μM was added and incubated for 30 minutes at 37 °C. Cells were washed with PBS and fixed with 3.7 % paraformaldehyde (PFA) for 15 min. The intracellular fluorescence of the SH-SY5Y cells was analyzed on a Leica TCS SP5 (Leica Microsystems, Wetzlar, Germany) with a HCX PL APO 63 × 1.4 oil immersion objective, under UV light by using a 488 nm excitation laser for CellROX and collecting the emission with a 515-560 nm filter range.

**Oligomer binding to cells.**

SH-SY5Y cells were seeded onto glass coverslips (Ibidi, Gräfelfing, Germany) and treated for 45 minutes with 10 μM of type B* oligomers or type B* pretreated for 15 minutes with an equimolar concentration of PSMα3. Cells were then washed with PBS and fixed with 3.7 % PFA for 15 minutes. Then cells were washed with PBS containing 0.1 % Triton X-100 for 10 min. Cells were blocked with 5 % BSA-PBS and incubated with 1/200 dilution rabbit polyclonal anti-αS antibody (Abcam, Cambridge, UK) overnight at 4 °C, and with 1:1000 anti-rabbit secondary antibodies conjugated with AF488. Cell nuclei was stained using Hoescht 33342 at a concentration of 0.5 μg/mL for 5 min. Images of intracellular αS were obtained under UV light using double excitation at 488 nm and 350 nm lasers, for AF488 and Hoescht, and the emission was collected at 515-560 nm and 405 nm, respectively.

**Redesign of PSMα3 variants**

To guide and assist the design of PSMα3 peptide variants some computational tools were employed. Briefly, AGADIR was used to predict the helical propensity of the peptide variants based on the helix/coil transition theory[8]. FoldX allows a rapid evaluation of the effect of mutations on the stability, folding and dynamics of proteins[9]. We exploited it to evaluate if the designed mutations may compromise the stability of the α-helix specially regarding extensive redesign or those involving electrostatic repulsions. The peptides mean hydrophobicity (*H*), and their helical hydrophobic moment ($\mu_H$), a measure of the amphiphilicity of a helix, were calculated according to Eisenberg and coworkers[10].

**Statistical analysis**

Statistical analyses were performed using Prism (GraphPad Software Inc., La Jolla, CA, USA), and mean values were compared using unpaired two-tailed t tests (Welch-corrected). No samples, or data points were excluded from the reported analyses.

**References**

1. Pujols, J. et al. Small molecule inhibits alpha-synuclein aggregation, disrupts amyloid fibrils, and prevents degeneration of dopaminergic neurons. *Proc Natl Acad Sci U S A* **115**, 10481-10486 (2018).

2. Pena-Diaz, S. et al. ZPD-2, a Small Compound That Inhibits alpha-Synuclein Amyloid Aggregation and Its Seeded Polymerization. *Front Mol Neurosci* **12** (2019).

3. Cascella, R. et al. Probing the Origin of the Toxicity of Oligomeric Aggregates of alpha-Synuclein with Antibodies. *ACS Chem Biol* **14**, 1352-1362, doi:10.1021/acschembio.9b00312 (2019).

4. Cremades, N. et al. Direct observation of the interconversion of normal and toxic forms of alpha-synuclein. *Cell* **149**, 1048-1059 (2012).

5. Fusco, G. et al. Structural basis of membrane disruption and cellular toxicity by alpha-synuclein oligomers. *Science* **358**, 1440-1443 (2017).

6. Chen, S. W. & Cremades, N. Preparation of alpha-Synuclein Amyloid Assemblies for Toxicity Experiments. *Methods Mol Biol* **1779**, 45-60 (2018).

7. Kruger, D., Ebenhan, J., Werner, S. & Bacia, K. Measuring Protein Binding to Lipid Vesicles by Fluorescence Cross-Correlation Spectroscopy. *Biophys J* **113**, 1311-1320 (2017).

8. Munoz, V. & Serrano, L. Elucidating the folding problem of helical peptides using empirical parameters. *Nat Struct Biol* **1**, 399-409 (1994).

9. Schymkowitz, J. et al. The FoldX web server: an online force field. *Nucleic Acids Res* **33**, W382-388 (2005).

10. Eisenberg, D., Weiss, R. M. & Terwilliger, T. C. The helical hydrophobic moment: a measure of the amphiphilicity of a helix. *Nature* **299**, 371-374 (1982).
